# Cerebral dopamine neurotrophic factor (CDNF) reduces myocardial ischemia/reperfusion injuries by the activation of PI3K-AKT via KDEL-receptor binding

**DOI:** 10.1101/683110

**Authors:** Leonardo Maciel, Dahienne Ferreira de Oliveira, Fernanda Mesquita, Hercules Antônio da Silva Souza, Leandro Oliveira, Fernando L. Palhano, Antônio Carlos Campos de Carvalho, José Hamilton Matheus Nascimento, Debora Foguel

## Abstract

CDNF (Cerebral Dopamine Neurotrophic Factor) belongs to a new family of NF, which presents several beneficial activities beyond the brain. Little is known about CDNF in the cardiac context. Herein we investigate CDNF effects in cardiomyocytes under endoplasmic reticulum (ER)-stress and in whole rat hearts subjected to ischemia/reperfusion (I/R). We showed that CDNF is secreted by cardiomyocytes stressed by thapsigargin and by isolated hearts subjected to I/R. CDNF protects human and mouse cardiomyocytes against ER-stress by restoring the calcium transient, and isolated heart against I/R injuries by reducing the infarct area and avoiding mitochondrial impairment. This protection is abrogated by wortmannin (PI3K-inhibitor) or by heptapeptides containing KDEL sequence, which block KDEL-receptor. These data suggest that CDNF induces cardioprotection via KDEL receptor binding and PI3K/AKT activation. This is the first study to propose CDNF as a cardiomyokine and to unravel the receptor and signaling pathway for this interesting family of NF.

## Introduction

CDNF (cerebral dopamine neurotrophic factor) together with MANF (mesencephalic astrocyte derived neurotrophic factor) form a new family of neurotrophic factors (NF) that are structurally unrelated to the other three families of NF (Lindahl et al., 2017). Human CDNF shares 59% amino-acid identity with human MANF and both have signal peptides that address them to the endoplasmic/sarcoplasmatic reticulum (ER/SR) (Liu et al., 2015; Lindahl et al., 2017).

Our group was the first to solve by nuclear magnetic resonance spectroscopy the structure of full-length human CDNF in solution, revealing a protein with two folded domains, the N- and the C-domains, with perfect superposition to the MANF structure (Latge et al., 2015). Several studies have elucidated the striking effects of MANF and CDNF in protecting neuronal cells against several injuries (reviewed in Lindahl et al., 2017), specially the dopaminergic neurons (Lindholm et al., 2007; Latge et al., 2015), thereby identifying these NFs as candidates for PD therapy (Voutilainen et al., 2011; Bäck et al., 2013; Voutilainen et al., 2015; Sampaio et al., 2017).

Although the precise mechanism(s) of action of MANF and CDNF is not completely understood, ER/SR stress relief seems to be one of their main activities. Several studies have shown that MANF and CDNF are induced by ER/SR stress (Apostolou et al., 2008; Tadimalla et al., 2008; Glembotski et al., 2012; Hartley et al., 2013; Liu et al., 2015). ER/SR stress has been implicated in a series of insults, including reduction in ER/SR calcium stores, oxidative stress, and an altered protein glycosylation pattern, among others. All of these cause the accumulation of misfolded proteins in the ER/SR, triggering a series of cellular events, collectively called the unfolding protein response (UPR). UPR activates the translation of a set of very specific proteins, which relieve the stress and, if necessary, induce cell apoptosis (Hetz, 2012; Lee et al., 2013;).

ER/SR stress has been implicated in human pathologies such as neurodegenerative and cardiac diseases, diabetes etc. Different from other proteins involved in ER/SR stress, MANF and CDNF are secreted to the extracellular milieu where they exert autocrine/paracrine/endocrine effects (Sun et al., 2011; Glembotski et al., 2012; Oh-Hashi et al., 2012; Liu et al., 2018). Interestingly, MANF and CDNF have in their C-terminal portion the sequences RTDL or KTEL, respectively. These sequences closely resemble the classical ER/SR retention signal, KDEL (Lys-Asp-Glu-Leu), which has a high affinity for KDEL-receptors (KDEL-R). This sequence is present in several ER/SR-resident proteins such as GRP78 (glucose-regulated protein 78), calreticulin and PDI (protein disulfide isomerase) (Raykhel et al., 2007; Capitani et al., 2009), for instance. The KDEL-R is involved in the retrieval of proteins from Golgi back to ER/SR, avoiding their secretion. The presence of degenerated KDEL sequences in MANF and CDNF suggests that their affinity for KDEL-R is diminished, which contributes to their increased secretion (Glembotski et al., 2012). Indeed, deletion of the RTDL sequence from MANF enhances its secretion in mouse mesencephalic astrocytes, suggesting that MANF-KDEL-R interaction in the ER/SR compartment regulates MANF secretion (Oh-Hashi et al., 2012).

The cardioprotective activity of MANF has been investigated in detail (Tadimalla et al., 2008; Glembotski et al., 2011; Glembotski et al., 2012), but even though the heart and skeletal muscle express relatively high amounts of CDNF (Lindholm et al., 2007), there is only one report in the literature showing increased levels of CDNF expression in ER/SR stressed cells. Tunicamycin (TN), an ER/SR stressor, raised the expression of CDNF in H9C2 cells, a rat cardiomyoblast cell line, while a CDNF-hemagglutinin construct enhanced resistance of cardiomyocytes to TN, reducing apoptosis (Liu et al., 2018). However, the authors investigated neither the mechanism of action of CDNF nor its role to cardiac functions.

The main goal of the present study was to investigate CDNF effects in cardiomyocytes. By using isolated hearts from rats and cell culture of cardiomyocytes from mice as well as human induced pluripotent stem cells differentiated into cardiomyocytes (hiPSC-dCM), we show that CDNF is secreted by cardiomyocytes when calcium stores are disturbed by thapsigargin (TG), an ER stressor, and in isolated hearts subjected to ischemia/reperfusion (I/R). Exogenously added CDNF protects the heart against the injuries induced by I/R or TG and this protection is abrogated in the presence of wortmannin, a PI3K-specific inhibitor, or by KDEL-containing heptapeptides, suggesting that the PI3K/AKT pathway is involved in CDNF-induced cardioprotection and the KDEL-R is a receptor for CDNF in the cardiac context.

## Results

### Thapsigargin (TG) induces ER stress and increases CDNF secretion in neonatal ventricular cardiomyocytes and in human induced pluripotent stem cells differentiated into cardiomyocytes (hiPSC-dCM)

Human iPSC-dCM cells were produced as described previously (Lian et al., 2013) and their differentiation was confirmed by expression of troponin T (TnT). Approximately 61% of TnT-positive cells were observed in our differentiation protocol (not shown). As seen in **Figure 1**, in the absence of TG, CDNF was found mainly inside the cells (**panels A and C, controls**) and very little of this NF appeared in the cell media (**panels B and D, controls**). However, after 20h in the presence of TG, there was an increase in the amount of CDNF detected in the cell lysates (**panels A and C, TG**), and considerable amounts were also detected in the cell media (**panels B and D, TG**). As seen in **Figure 1E**, treatment with TG (1μmol/L/20h) significantly increased the expression of GRP78 to 1.7-fold when compared to controls, confirming that these cells were under ER/SR stress condition. Interestingly, when the cells were pre-treated with 1μmol/L exogenous CDNF (exoCDNF) for 15min before TG addition, there was no increase in GRP78 (**panel E**), a marker of ER stress, suggesting that exoCDNF is preventing TG-induced ER/SR stress. **Figure 1G** shows that isolated hearts subjected to I/R had their levels of CDNF increased to 27 times the control values, and this was accompanied by an increase in CHOP (CCAAT/-enhancer-binding protein homologous protein), another ER/SR stress marker, to ~1.7-fold, suggesting that these hearts were under ER/SR stress. ExoCDNF added before I/R led to a significant decrease in CHOP levels by approximately 50% (**Figure 1F**).

**Figure 1:**
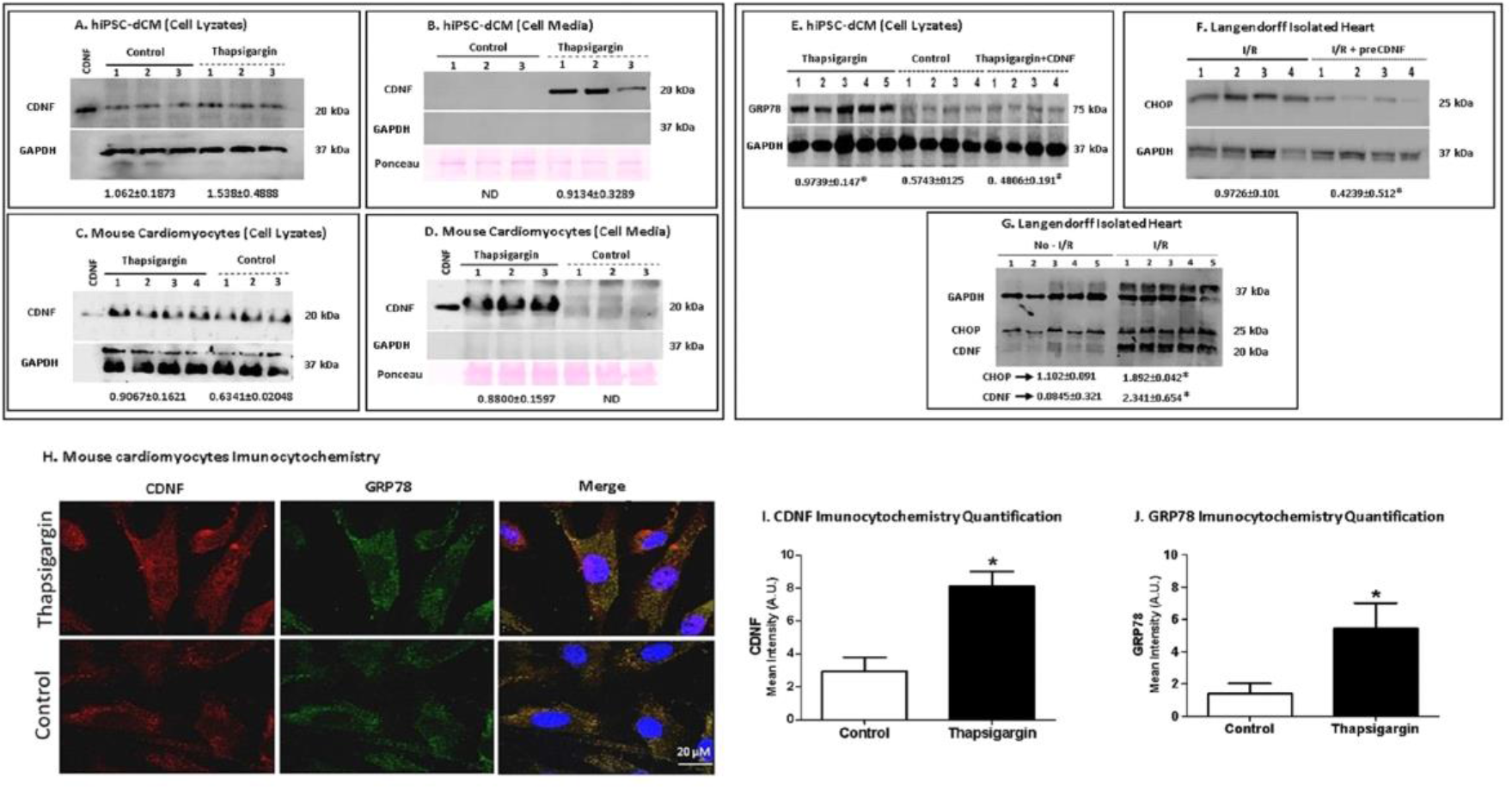
Thapsigargin (TG) treatment increases the levels of CDNF in human and mouse cardiomyocytes as well as its secretion to the extracellular media. Human iPSC-dCM (**A, B**) or mouse cardiomyocytes (**C, D**) were treated for 20h with TG (1μmol/L). **CDNF blocks the TG- and I/R-induced increase in the levels of GRP78 and CHOP in hearts and human cardiomyocytes**. (**E**) Levels of GRP78 in cell extracts of hiPSC-dCM cultures treated with TG (1μmol/L) or with CDNF (1μmol/L) prior to TG. (**F**) Levels of CHOP and CDNF from rat hearts subjected to I/R protocol. (**G**) Levels of CHOP from hearts subjected to I/R after CDNF treatment (1μmol/L/5min). (**H**) Levels of CDNF and GRP78 in mouse cardiomyocytes before and after TG treatment, measured by confocal imaging. (**I** and **J**) Immunocytochemistry quantification of CDNF and GRP78. The levels of CDNF, GRP78 and CHOP are normalized to GAPDH levels in cell lysate and to Ponceau red in extracellular media. Recombinant CDNF was used as control. Each lane represents a different cell culture and the quantifications of protein expression (± S.E.M.) are shown below each blot. *P<0.05 *vs*. control and #P<0.05 *vs*. TG. ND, not detected.

Taken together these data suggest that TG and I/R, two ER/SR stressors, increase the concentration of CDNF, GRP78 and CHOP in hiPSC-dCM and neonatal mice cardiomyocytes as well as in the intact heart. Pre-treatment of the cells or the hearts with exoCDNF blocked the increase in the two UPR markers, GRP78 and CHOP, suggesting that CDNF is an anti-ER/SR stressor in the cardiac context. Interestingly, TG treatment, in addition to increasing the cellular levels of CDNF in mouse or human cardiomyocytes, also increases its secretion to the extracellular milieu, which is consistent with a protective activity of this NF.

### ExoCDNF prevents the Ca^2+^-transient depletion induced by thapsigargin

As seen in **Figure 2**, TG treatment (1μmol/L/20h) reduced calcium transients of hiPSC-dCM (reduced spike amplitudes in **panel A**, quantified in **panel B**). However, pre-incubation of these cells with CDNF (1μmol/L/1h) upon TG addition potentiated the calcium transients to values even greater than the control, suggesting that exoCDNF exerts a direct effect on cardiomyocyte calcium homeostasis under stress conditions, probably by improving calcium uptake by the ER/SR, an outcome that may prevent cellular injuries provoked by cytosolic calcium overload (**Figure 2**). The addition of wortmannin abrogates the effects of CDNF on calcium transients after TG addition (to be described later).

**Figure 2:**
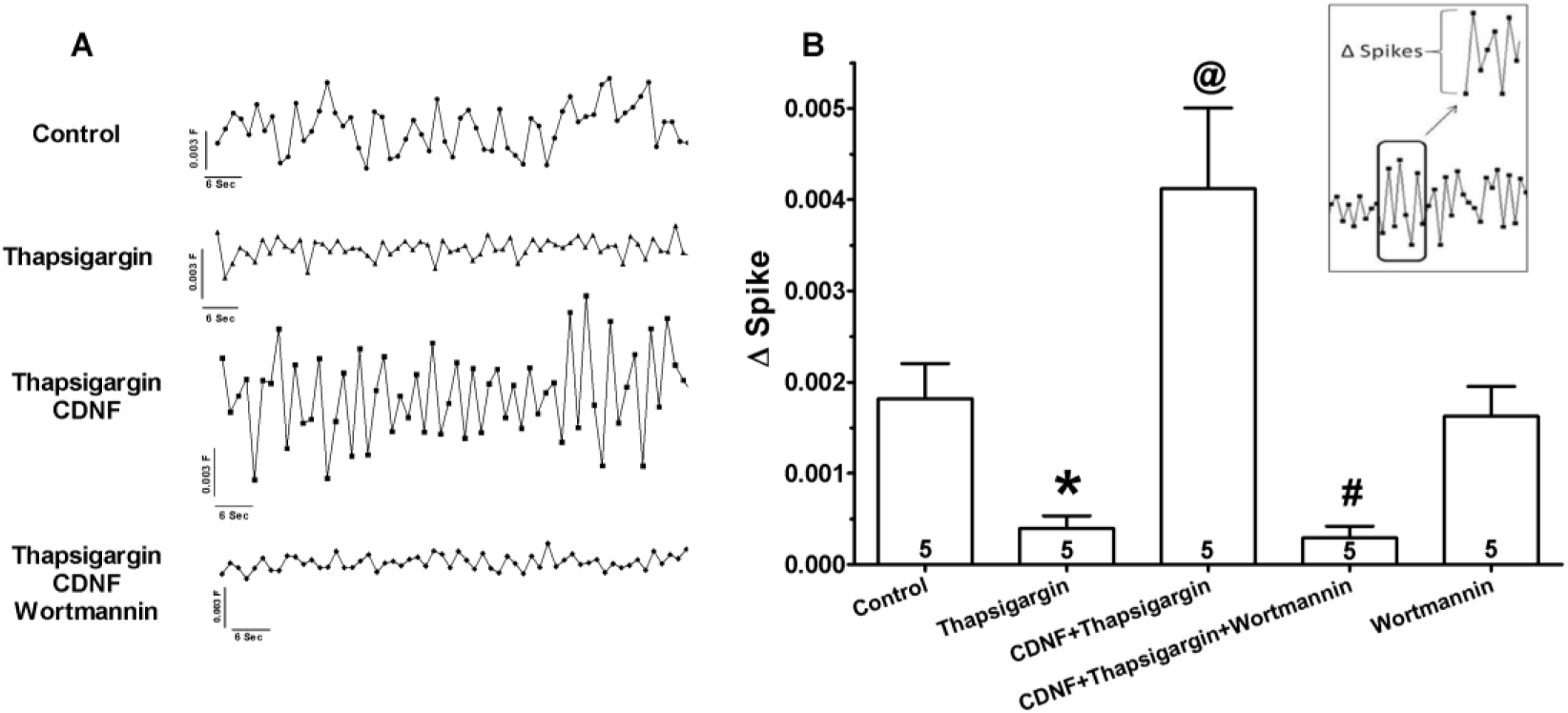
ExoCDNF restores the calcium transient in hiPSC-dCM perturbed by TG treatment, and wortmannin abrogates this beneficial effect of CDNF. Groups: Control, TG, TG+CDNF and TG+CDNF+wortmannin **(representative traces in panel A)**. (**B**) ΔSpike (average of 20 peaks for each experiment) represents the difference between the calcium level in a specific top peak in relation to the previous lower peak (see the inset) Data are means ± S.E.M. Number inside each column is *n* of hiPSC-dCM cultures tested. ^*^P<0.01 *vs*. control; ^@^P<0.01 *vs*. TG and ^#^P<0.01 *vs*. CDNF+TG.

### Cardioprotective effects of exoCDNF on isolated hearts

Initially, we evaluated whether rat heart chambers could express endogenous CDNF in significant amounts. As shown in **Figure 3** this does occur with the ventricles expressing greater amounts of CDNF per unit of total protein than the atria.

**Figure 3:**
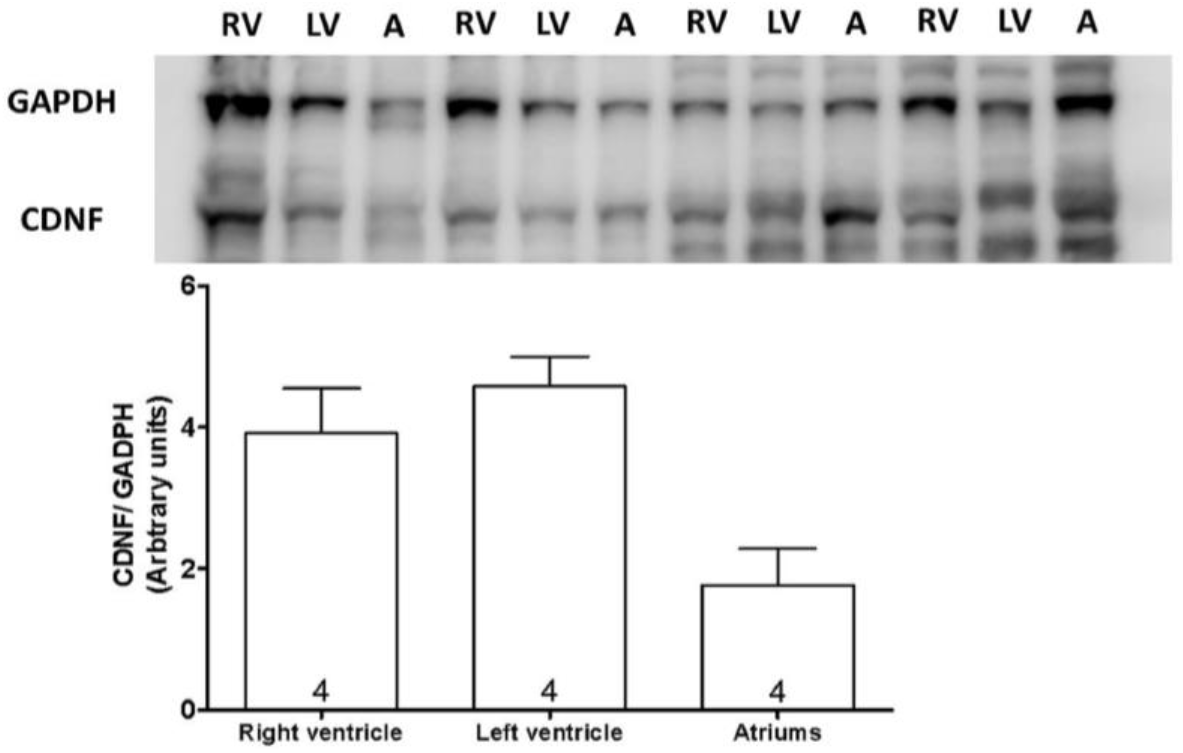
Endogenous CDNF expression in rat heart chambers. Extracts from different heart compartments (50μg protein) were fractionated by SDS-PAGE followed by immunoblotting for CDNF and GAPDH. The levels of CDNF were estimated by normalizing the intensity of the CDNF band to the GAPDH band as shown in the graph. Data are means ± S.E.M. Number in each column is *n* of hearts. RV= right ventricle; LV = left ventricle; A = atria. Fresh untreated hearts were used in these experiments.

Next, the cardioprotective activity of CDNF was evaluated by perfusion with exoCDNF as a preconditioning treatment (**Figure 4 A and B**), and as a postconditioning treatment (**Figure 4 C and D**). Since there was no significant difference in the baseline of the left ventricular developed pressure (LVDP) among the groups (**Table 1**), all LVDP changes are expressed as percentage of the baseline values, which were taken as 100%. Under ischemic condition all control hearts exhibited a rapid reduction of LVDP down to zero, recovering poorly (~20%) during reperfusion (**panels A and C, circles**). Interestingly, preCDNF and postCDNF treatments enhanced these recoveries considerably, up to 70% and 50%, respectively (**panels A and C, squares**). Regarding left ventricular end-diastolic pressure (LVEDP; **panels B and D**), preCDNF kept the pressure low during perfusion (40 mmHg, **squares**) in relation to the control, which rose to ~70 mmHg at the end of the reperfusion process (**circles**). However, postCDNF treatment presented no significant alteration in LVEDP compared to control (**panel D**).

**Figure 4:**
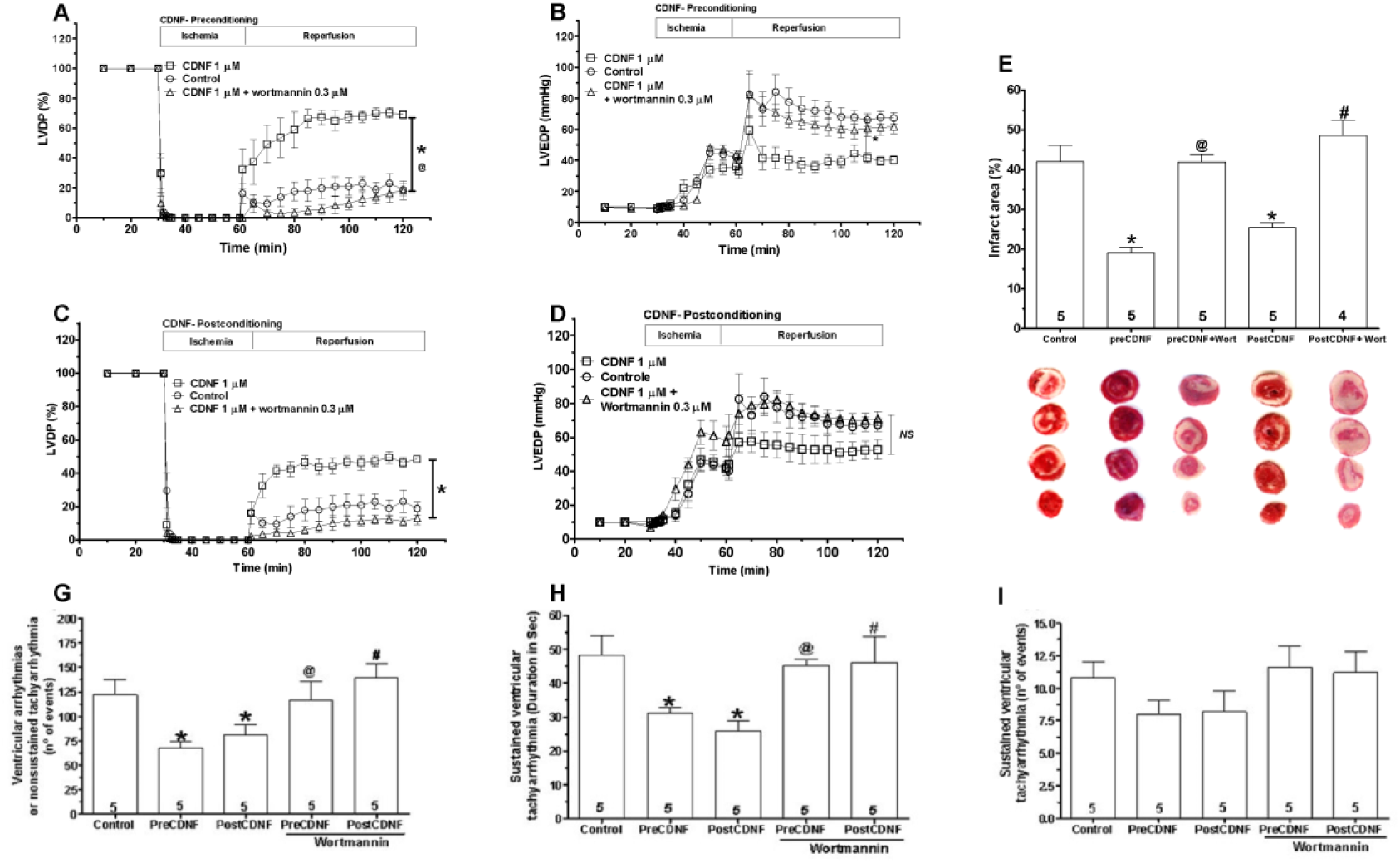
ExoCDNF protects intact rat hearts from injuries provoked by I/R and protection is prevented by wortmannin, a PI3K/AKT antagonist. Time courses of left ventricular developed pressure (LVDP) (**A, C**) and left ventricular end-diastolic pressure (LVEDP) (**B, D**) during I/R protocol. In **A** and **B**, CDNF (1μmol/L) was perfused 5min before ischemia (preconditioning) and in **C** and **D**, during the first 5min of reperfusion (postconditioning). Wortmannin (0.3μmol/L) was perfused 5min before CDNF. (**E**) **Infarct area of hearts**. Representative cross-section images of TCC-stained ventricle hearts subjected to I/R. (**F**) Number of ventricular arrhythmia or nonsustained tachyarrhythmia events or (**G**) Number of events and (**H**) the duration of sustained ventricular tachyarrhythmia produced during reperfusion. Values are expressed as means ± S.E.M. Number in each column is *n* of hearts. * P<0.01 *vs* control; ^@^P<0.01 *vs* preCDNF and ^#^P <0.01 vs postCDNF.

**Table 1:** Left ventricular developed pressure of isolated perfused rat hearts in mmHg.

| Groups | Time | Rat hearts LVDP [mmHg] |  |  |
| --- | --- | --- | --- | --- |
| Control (n=5) | baseline | 103.1 | ± | 8.9 |
| preCDNF (n=5) | baseline | 97.9 | ± | 14.3 |
|  | after intervention | 102.5 | ± | 11.2 |
| **postCDNF(n=5)** | **baseline** | **101.9** | **±** | **18.2** |
| **preCDNF**<br>**+Wortmannin (n=5)** | **baseline** | **95.7** | **±** | **8.3** |
|  | **after intervention** | **88.3** | **±** | **10.2** |
| **postCDNF**<br>**+Wortmannin (n=5)** | **baseline** | **111.5** | **±** | **10.7** |
| **preCDNF+**<br>**TRPQTEL(n=5)** | **baseline** | **102.3** | **±** | **15.1** |
|  | **after intervention** | **96.1** | **±** | **7.2** |
| **preCDNF+**<br>**THPKTEL(n=5)** | **baseline** | **97.6** | **±** | **8.1** |
|  | **after intervention** | **91.7** | **±** | **13.4** |
| **preCDNF+**<br>**DRATSAL(n=5)** | **baseline** | **99.3** | **±** | **13.2** |
|  | **after intervention** | **102.4** | **±** | **6.9** |
| **PreCDNF+**<br>**αβ-CDNF(n=5)** | **baseline** | **107.4** | **±** | **10.4** |
|  | **after intervention** | **98.3** | **±** | **8.2** |
| **No-I/R (n=9)**<br>**(mitochondria assay)** | **baseline** | **101.6** | **±** | **6.9** |
| **I/R (n=9)**<br>**(mitochondria assay)** | **baseline** | **104.2** | **±** | **8.8** |
|  | **ischemia 5 min** | **0.0** | **±** | **0.0** |
|  | **ischemia 30 min** | **0.0** | **±** | **0.0** |
|  | **reperfusion 10 min** | **14.1** | **±** | **16.3** |
| **preCDNF (n=5)**<br>**(mitochondria assay)** | **baseline** | **110.8** | **±** | **7.5** |
|  | **after intervention** | **105.3** | **±** | **15.9** |
|  | **ischemia 5 min** | **0.0** | **±** | **0.0** |
|  | **ischemia 30 min** | **0.0** | **±** | **0.0** |
|  | **reperfusion 10 min** | **43.9** | **±** | **7.6 \*** |
| **PostCDNF (n=4)**<br>**(mitochondria assay)** | **baseline** | **99.8** | **±** | **10.9** |
|  | **after intervention** | **100.2** | **±** | **8.3** |
|  | **ischemia 5 min** | **0.0** | **±** | **0.0** |
|  | **ischemia 30 min** | **0.0** | **±** | **0.0** |
|  | **reperfusion 10 min** | **33.7** | **±** | **11.4** |
| **preCDNF**<br>**+Wortmannin (n=5)**<br>**(mitochondria assay)** | **baseline** | **113.3** | **±** | **11.6** |
|  | **after intervention** | **102.0** | **±** | **6.1** |
|  | **ischemia 5 min** | **0.0** | **±** | **0.0** |
|  | **ischemia 30 min** | **0.0** | **±** | **0.0** |
|  | **reperfusion 10 min** | **15.9** | **±** | **14.3 &** |
| **postCDNF**<br>**+Wortmannin (n=3)**<br>**(mitochondria assay)** | **baseline** | **102.1** | **±** | **19.7** |
|  | **after intervention** | **93.5** | **±** | **15.4** |
|  | **ischemia 5 min** | **0.0** | **±** | **0.0** |
|  | **ischemia 30 min** | **0.0** | **±** | **0.0** |
| | **reperfusion 10 min** | **11.3** | **±** | **12.5 \$** |

Mean of maximal developed left ventricular pressure (LVDP) of isolated perfused rat hearts. The LVDP was analyzed at different time points: at baseline, after the intervention, at 5 and 30 min of ischemia, and at 10 min of reperfusion, respectively. Mean ± SEM. \* P< 0.05 vs I/R, & P< 0.05 vs preCDNF, \$ P< 0.05 vs postCDNF.

Myocardial infarct areas evaluated after 60 min of reperfusion were measured in control, preCDNF and postCDNF hearts (**Figure 4 E**). As seen, while in the control hearts the mean infarct area reached 42%, the hearts subjected to preCDNF or postCDNF treatments showed substantial reduction in the infarct area to 19% and 25%, respectively.

As shown in **Figure 4 (panels F-H)**, there was a significant reduction in the mean number of ventricular arrhythmias or non-sustained tachyarrhythmia events (**panel F**) produced during the 60 min of reperfusion in preCDNF and postCDNF protocols. The number of sustained ventricular tachyarrhythmias was not significantly reduced by preCDNF or postCNDF treatments (**panel H**), although the duration of these tachyarrhythmias was reduced from ~48 s to 33 and 25 s in preCDNF and postCDNF, respectively (**panel G**).

### PI3K/AKT pathway is mediating the cardioprotective effects of exoCDNF

Next, we examined the signaling pathway involved in the cardioprotective effect conferred by exoCDNF. Surprisingly, only wortmannin was able to abrogate the cardioprotective effect of exoCDNF treatment (**Figure 4, triangles**), while the other inhibitors AG490 (JAK-STAT3 inhibitor), rottlerin and chelerythrine (PKC inhibitors) failed as displayed in **Figure 5**. As seen in **Figure 4**, the massive recovery in LVDP (**panel A**) or the maintenance of the low values of LEVDP (**panel B**) observed with preCDNF treatment before I/R injury (**squares**) were completely abolished in the presence of wortmannin (**triangles**) and values were similar to those of the control groups (**circles**). Because only wortmannin abrogated CDNF-induced cardioprotection in preCDNF condition, only this inhibitor was evaluated in postCDNF treatment, and again wortmannin abrogated LVDP recovery (**Figure 4C**). The pre-treatment with wortmannin led to an increase of infarct area in preCDNF and postCDNF to ~40% and ~45%, respectively, similar to the control (**Figure 4E**). The lack of effect of the other inhibitors on infarct area are depicted in **Figure 5**. In addition, treatment with wortmannin in preCDNF and postCDNF protocols abolished the reduction of ventricular arrhythmias in hearts subjected to I/R (**Figure 4 F and G**). Besides, as shown before in **Figure 2**, addition of wortmannin also abrogates the protective effects of CDNF on calcium transients after TG addition.

**Figure 5:**
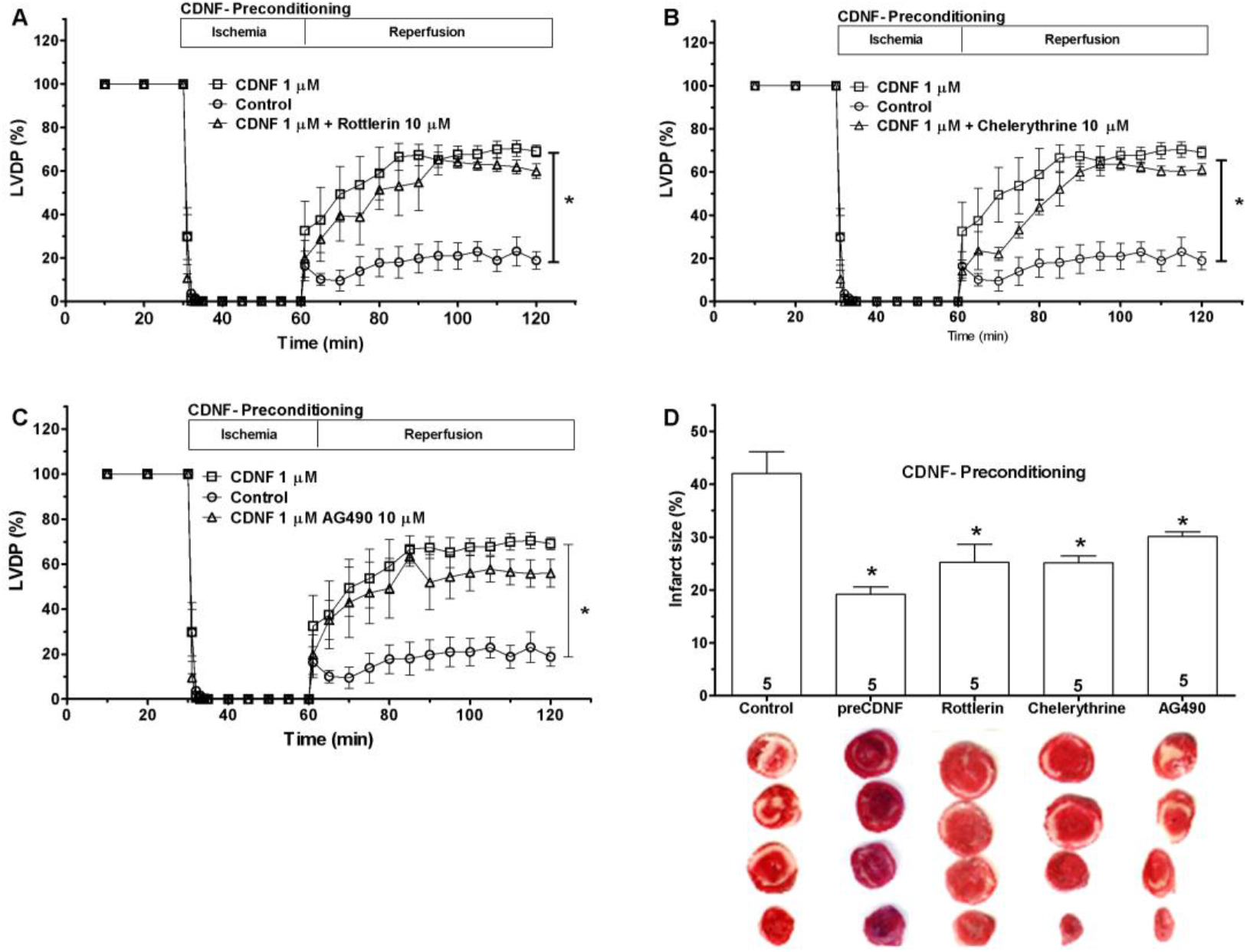
The cardioprotective activity of CDNF is not prevented by rottlerin (A), chelerythrine (B) or AG490 (C). Time courses of left ventricular developed pressure (LVDP) during I/R protocol (30min of global ischemia and 60 min of reperfusion) or when the hearts were subjected to a previous perfusion with CDNF (1 μmol/L/5 min - preconditioning) or with CDNF+inhibitor (5min before I/R). Controls (**circles**), CDNF treatment (**squares**) and CDNF+inhibitor (**triangles**). **(D) Rottlerin, chelerythrine and AG490 do not counteract the protective effect of CDNF in reducing the infarct area of hearts subjected to I/R**. Representative cross-sections of TCC-stained ventricles and quantification of control hearts subjected to I/R only, or I/R after preconditioning with CDNF or with CDNF and inhibitors. Numbers inside each bar are the number of hearts used. Data are means ± S.E.M. ^*^P < 0.001 *vs*. control.

The levels of phosphorylated AKT (p-Akt) were evaluated in hiPSC-dCM, mouse cardiomyocytes and isolated hearts before and after addition of exoCDNF (**Figure 6, panels A, B and C**, respectively). In all three models, the levels of p-Akt more than doubled after CDNF addition, except when CDNF was added after incubation with wortmannin.

**Figure 6.**
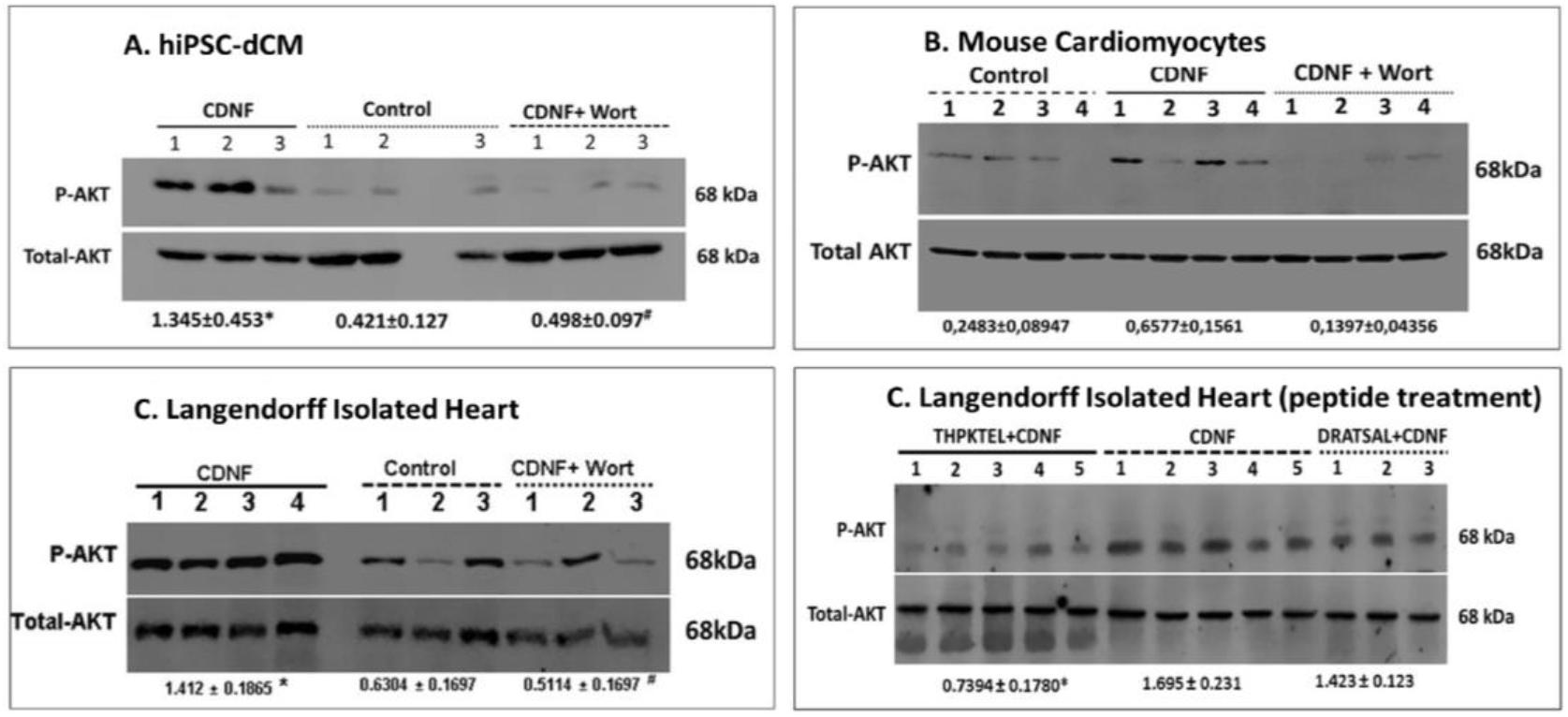
ExoCDNF treatment increases the level of phosphorylated AKT (p-Akt) in rat isolated hearts and in mouse and human cardiomyocytes. Cultures of (**A**) hiPSC-dCM or (**B**) mouse cardiac myocytes were treated with CDNF (1μmol/L/20h) or with wortmannin (0.3μmol/L/15min) before CDNF addition. In (**C**), rat isolated hearts were perfused with CDNF (1□mol/L/5min, preconditioning) before I/R either alone or in combination with wortmannin (0.3μmol/L/5min) before CDNF treatment. The intensities of the p-Akt bands were normalized to total-AKT levels. (**D**) **The peptide THPKTEL that binds to the KDEL-R at the cell membrane blocks CDNF-induced PI3K/AKT activation**. Rat isolated hearts were perfused with CDNF (1μmol/L/5min, preconditioning) before I/R either alone or in combination with THPKTEL or with the scrambled peptide DRATSAL (peptides added 5min before CDNF treatment and during the 5min CDNF treatment). The number of experiments is equal to number of lanes. ^*^P<0.001 *vs*. control ^#^P<0.01 *vs*. CDNF.

### exoCDNF exerts a protective effect on mitochondrial function after heart I/R

Protection of mitochondrial function during I/R has been reported to be beneficial to cardiomyocytes (Heusch, 2015). In order to investigate whether CDNF could exert any effect on mitochondrial function and whether wortmannin could counteract these effects, mitochondria were isolated from hearts subjected to I/R in the Langendorff model with or without perfusion of CDNF (preCDNF or postCDNF) and CDNF+Wortmannin (**Figure 7**).

**Figure 7.**
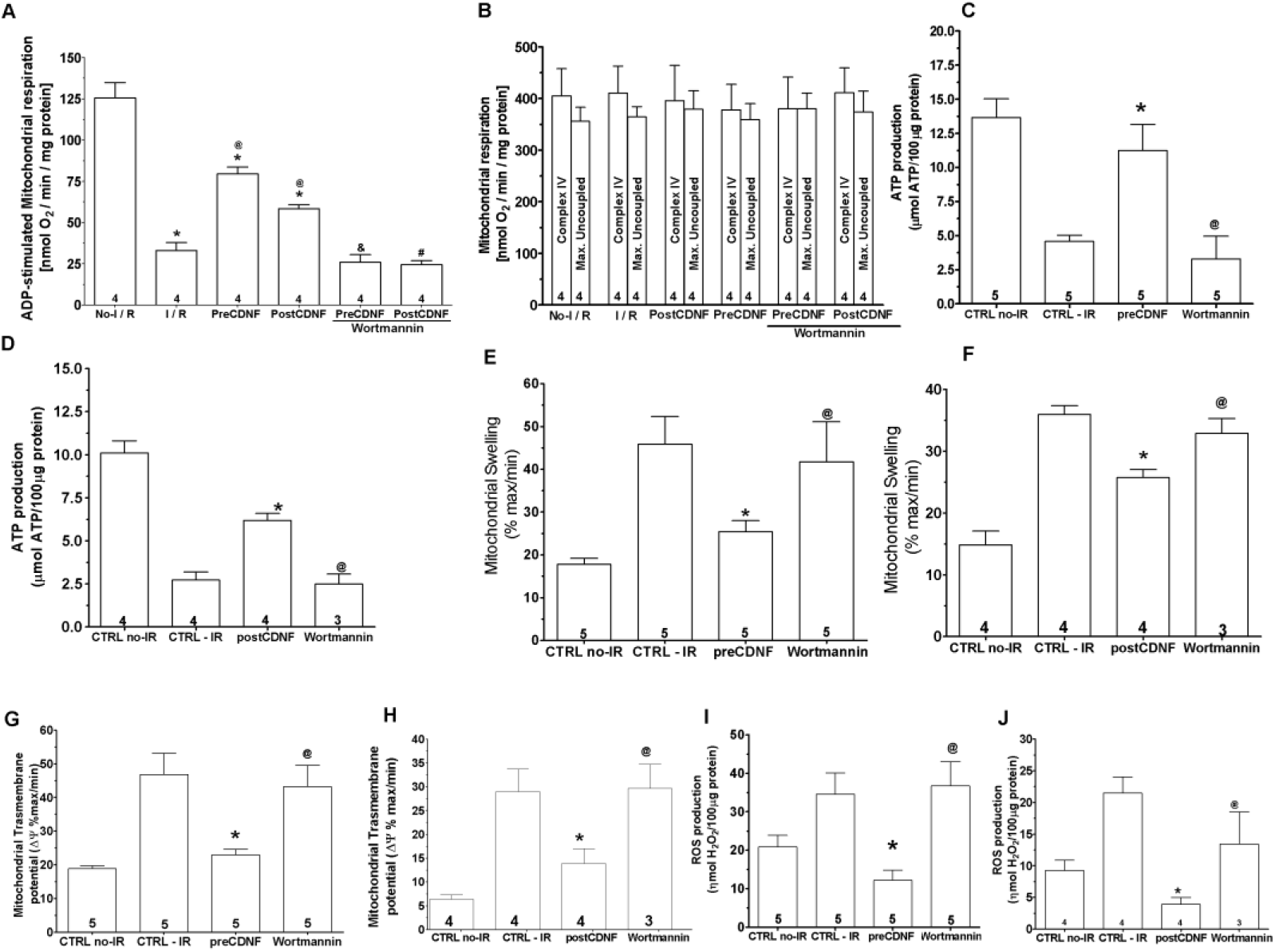
ExoCDNF reduces cardiac mitochondrial impairment induced by I/R and wortmannin abrogates the CDNF beneficial effects. (**A**) ADP-Stimulated complex I respiration; (**B**) Complex IV and maximal uncoupled oxygen respiration; (**C, D**) ATP production; **(E, F)** Mitochondrial swelling (**G, H**) Mitochondrial transmembrane potential (ΔΨ) and **(I, J)** Reactive oxygen species (ROS) production. Groups: No-I/R, I/R, preCDNF, postCDNF and preCDNF+Wortmannin and postCDNF+Wortmannin. In (**C-I**) preconditioning with CDNF or with CDNF+Wortmannin. In (**D-J**) postconditioning with CDNF and CDNF+Wortmannin. ^*^P<0.05 *vs*. I/R; ^@^P<0.05 *vs*. No-I/R; ^&^P<0.05 *vs*. preCDNF and ^#^P<0.05 *vs*. postCDNF. Number in each column is *n* of hearts.

As expected, ADP-stimulated complex I respiration was reduced in the I/R group and this decrease was restored partially, but significantly, by pre or postCDNF treatments (1μmol/L); but the pre-treatment was more effective (**panel A**). Wortmannin abrogated the improvement conferred by preCDNF and postCDNF. The same scenario was found for ATP production (**panels C and D**) and mitochondrial swelling (**panels E and F**), indicating that the I/R-induced oxidative phosphorylation impairment and the increased mitochondrial volume were partially reversed by pre or postCDNF treatments, acting through the PI3K/AKT pathway. Mitochondrial complex IV respiration and maximal oxygen uptake of uncoupled mitochondria were not different between groups, reflecting an equal loading of viable mitochondria (**panel B**). Mitochondrial transmembrane potential (Δψ) showed an increase due to I/R, reflecting the inner transmembrane depolarization. PreCDNF and postCDNF treatments ameliorated this dangerous Δψ increase and wortmannin abrogated the beneficial effects of CDNF (**panels G and H, respectively**). Next, mitochondrial ROS production was measured and, as expected, I/R led to an enhancement in ROS levels, but pre or postCDNF treatments reduced ROS only when wortmannin was absent (**panels I and J, respectively**).

To test whether exoCDNF could exert a direct mitochondrial protection independent of PI3K/AKT activation, mitochondria from non-ischemic hearts were isolated and subjected to hypoxia/reoxygenation with or without previous incubation with CDNF. As shown in **Figure 8A**, the ADP-stimulated complex I respiration was reduced in hypoxia/reoxygenation group compared to control group, but in this case CDNF was ineffective in blocking or reversing this effect, confirming that intact cells are necessary for the beneficial effects of CDNF. Mitochondrial complex IV respiration and maximal oxygen uptake of uncoupled mitochondria were not different between groups, reflecting an equal loading of viable mitochondria (**Figure 8B**).

**Figure 8:**
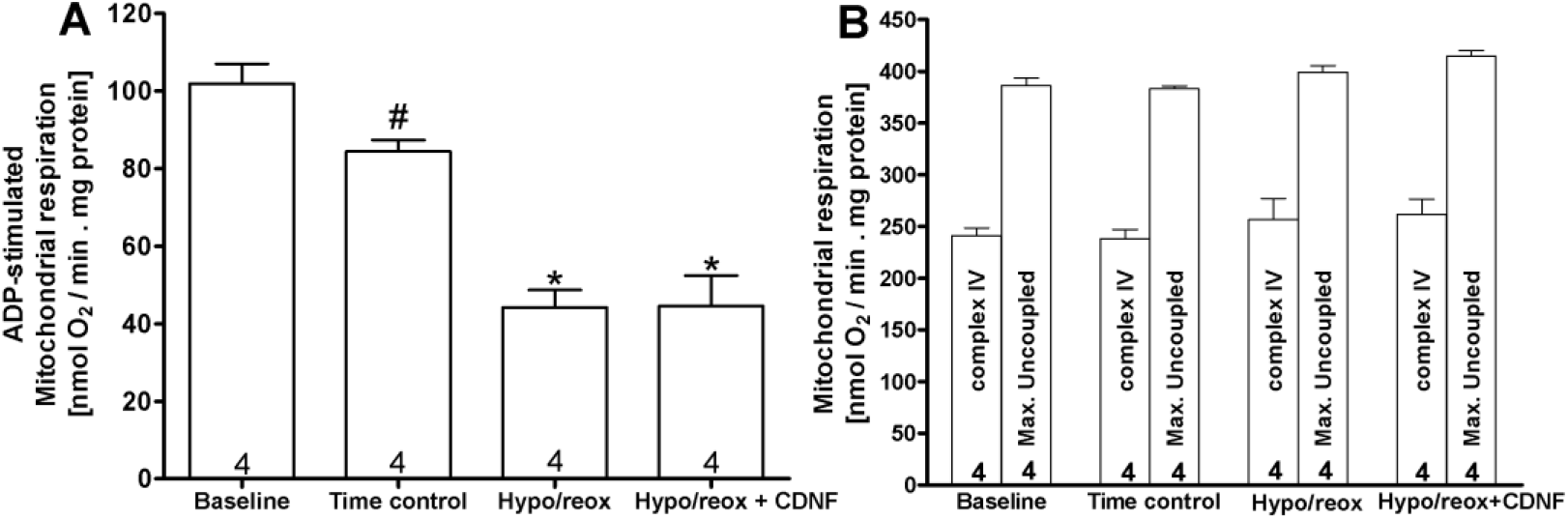
CDNF does not protect isolated mitochondria from hypoxia/reoxygenation. **(A)** ADP-Stimulated complex I respiration and (**B**) Complex IV-induced respiration with TMPD and ascorbate, compared to the maximal uncoupled oxygen uptake with FCCP present. The mitochondria were isolated from naive rat hearts and then subjected to hypoxia/reoxygenation in the absence or in the presence of CDNF (1 μmol/L). Groups: Baseline; Time control =10 min of mitochondria incubation in the chamber before the experiment; Hypo/reox=10min of hypoxia followed by reoxygenation; Hypo/reox+CDNF=CDNF incubation (1 μmol/L) before Hypo/reox. ^*^P<0.05 *vs*. time control; ^#^P<0.05 *vs*. baseline. Number in each bar is *n* of hearts.

### KDEL-R in the membrane binds CDNF and mediates its cardioprotective activity

We postulated that KDEL-R, which has been found in the cell membrane (Henderson et al., 2013; Becker et al., 2016; Ruggiero et al., 2017) and has on its operator sequence an ER stress-responsive element (ERSE) (Mizobuchi et al., 2007; Oh-Hashi et al., 2013), might function as a CDNF receptor. To investigate this, we perfused hearts before CDNF treatment with two heptapeptides, which incorporate the last seven amino-acid residues of human (THP**KTEL**) and rat (TRP**QTEL**) CDNF. The rationale was to block CDNF’s putative binding site on the KDEL-R, thus impairing CDNF function. As seen in **Figure 9**, (**TRPQTEL, panels A-C and THPKTEL, panels D-F**), both peptides were able to prevent the cardioprotection induced by preCDNF treatment, abolishing the beneficial recovery of LVDP (**panels A and D, diamonds**), the beneficial reduction of LVEDP (**panels B and E, diamonds**), and the decrease in infarct areas of the hearts (**panels C and F**) induced by CDNF. As seen in these panels, perfusion with the peptides alone was not able to confer cardioprotection (**triangles in all panels**), suggesting that the occupancy of the binding site on the KDEL-R by a ligand is not enough for full activation of the downstream pathway and cardioprotection.

**Figure 9:**
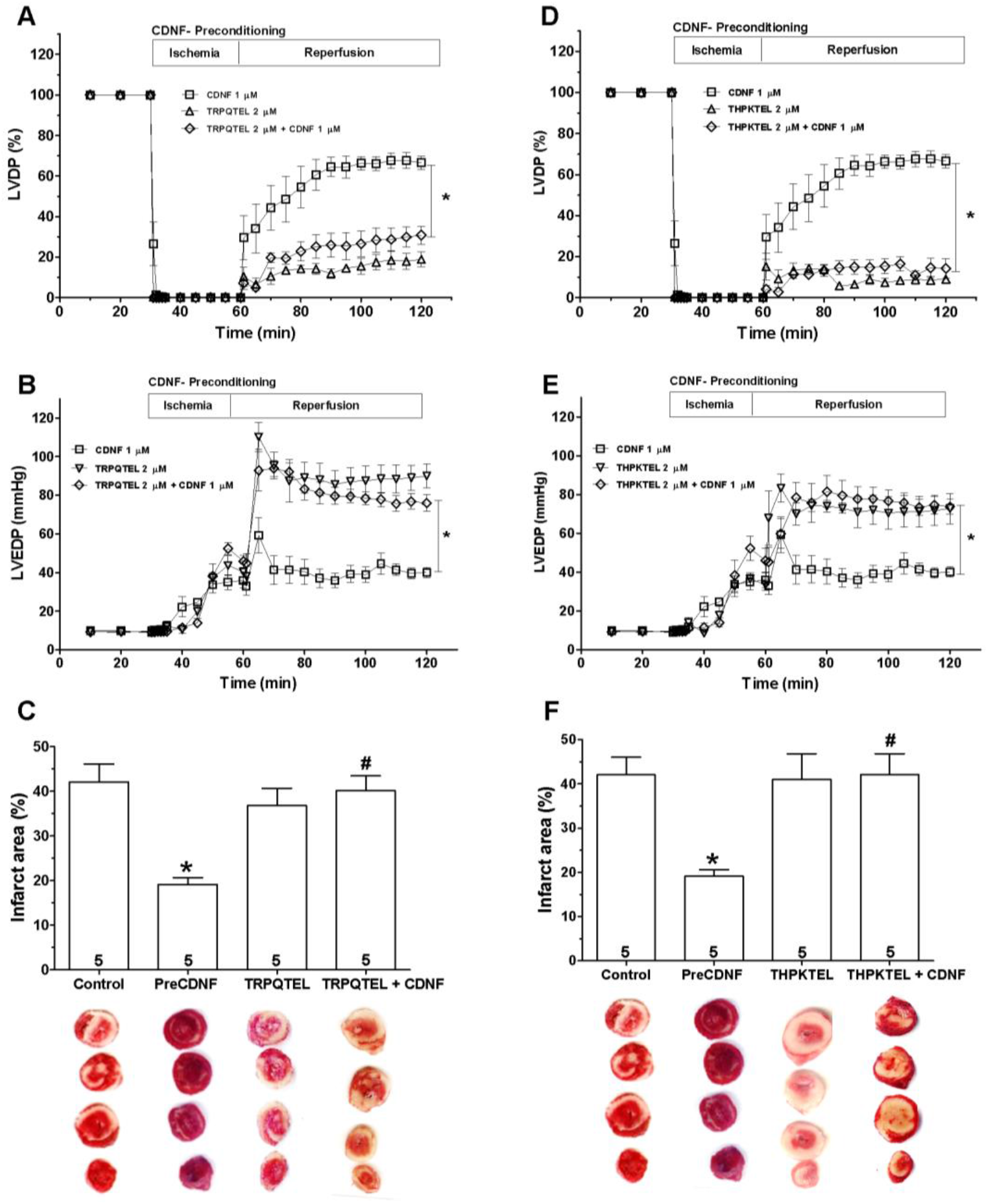
The cardioprotective effect of exoCDNF is blocked by heptapeptides that bind to KDEL-R. Time courses of left ventricular developed pressure (LVDP) (**A, D**) and left ventricular end-diastolic pressure (LVEDP) (**B, E**) during I/R protocol. CDNF (1μmol/L), peptide (in **A** and **B**, TRPQTEL and in **C** and **E**, THPKTEL; 2μmol/L of each peptide) or CDNF (1μmol/L)+peptide (2□mol/L) were perfused before ischemia (5min). **(C, F) infarct area of hearts**. Representative cross-section images of TCC-stained ventricles hearts subjected to I/R. Data are means ± S.E.M of 5 hearts. ^*^P<0.001 *vs*. control; ^#^P<0.001 *vs*. preCDNF.

**Figure 10** shows that the peptide **DRATSAL** used as a scrambled peptide by (Henderson et al., 2013) was not able to abolish the protective effect of CDNF in I/R experiments. Interestingly, THP**KTEL** was able to abolish CDNF-induced p-AKT enhancement, an effect that was not observed with the scrambled peptide. This result reinforces the link between KDEL-R activation and the PI3K/AKT signaling pathway (**Figure 6D**).

**Figure 10:**
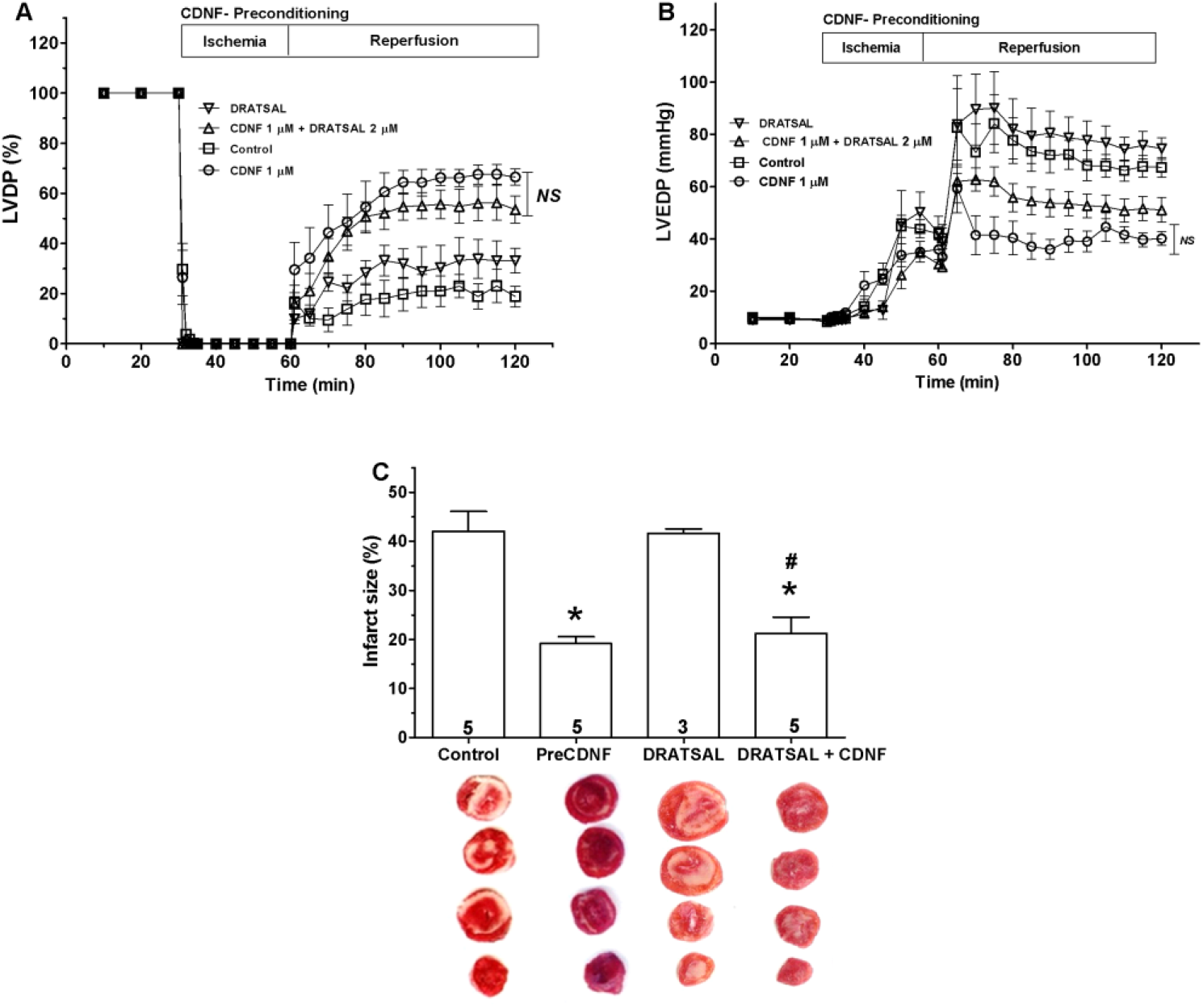
The cardioprotective effect of exoCDNF is not blocked by the scrambled peptide DRATSAL. Time course of (**A**) left ventricular developed pressure (LVDP) and (**B**) left ventricular end-diastolic pressure (LVEDP) during I/R protocol (30min of global ischemia and 60min of reperfusion). As indicated by the different tracings, CDNF (1 μmol/L), peptide (DRATSAL, 2μmol/L) or CDNF (1μmol/L)+DRATSAL (2μmol/L) were perfused before ischemia (5min). Control (**squares**), CDNF (**circles**), DRATSAL alone (**inverted triangles**) and CDNF+DRATSAL (**triangles**). **(C) DRATSAL did not block the decrease in infarct area induced by CDNF after I/R**. Representative cross-sections of TCC-stained ventricles and quantification of control hearts subjected to I/R or after preconditioning with CDNF, DRATSAL alone or CDNF+DRATSAL. Data are means ± S.E.M. The number of hearts used in each experiment is shown inside the bars. ^*^P < 0.001 *vs*. control; ^#^P < 0.001 *vs*. DRATSAL.

Finally, **Figure 11** shows that pre-incubation of CDNF with the anti-CDNF antibody (0.5μg/ml perfused for 5 min prior to the I/R protocol) completely abrogates the cardioprotective role (**panel A**, LVDP; **panel B**, LVEDP, and **panel C**, infarct area), probably because this antibody recognizes the C-terminal sequence of CDNF and prevents its binding to the KDEL-R.

**Figure 11:**
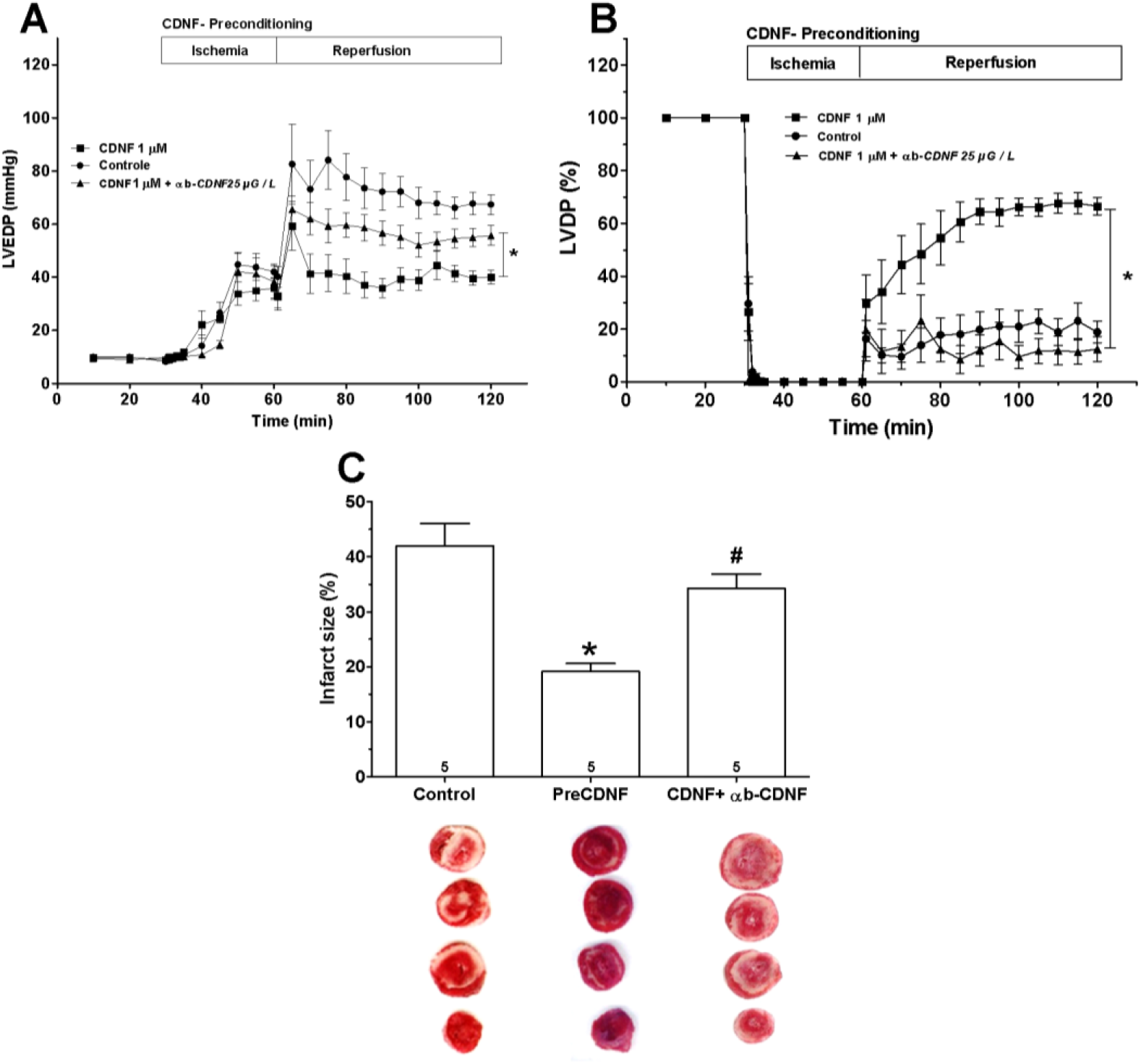
The cardioprotective effect of preconditioning with exoCDNF is blocked by an anti-CDNF antibody. Time courses of (**A**) left ventricular developed pressure (LVDP) and (**B**) left ventricular end-diastolic pressure (LVEDP) during I/R protocol and (**C**) Infarcted area. CDNF (1□mol/L) or CDNF+antibody were perfused before ischemia (5min). In **C**, Representative cross-section images of TCC-stained ventricles hearts subjected to I/R. Data are means ± S.E.M of 5 hearts. *^P^<0.001 *vs*. control; ^#P^<0.001 *vs*. preCDNF.

## Discussion

Cardiomyokines are proteins secreted by a healthy or a diseased heart displaying a beneficial autocrine/paracrine function (Glembotski et al., 2011; Doroudgar and Glembotski, 2011; Maciel et al., 2017). The present study is the first to characterize CDNF as an effective cardiomyokine that protects whole hearts, mice and especially human cardiomyocytes (iPS derived) in culture against injuries provoked by calcium overload, specifically, in our study, I/R and TG. As shown by others, MANF has cardioprotective activity (Tadimalla et al., 2008; Glembotski et al., 2012) and here we have extended this concept to CDNF, conferring on this member of this unusual family of neurotrophic factors a cardioprotective activity as well. Thus, CDNF and MANF can be considered as neuro/cardiokines and it is possible that other beneficial effects will be described in the near future for this interesting family of proteins. In fact, it has been shown that MANF and CDNF are also important regulators of inflammation and tissue repair (Neves et al., 2016; Sousa-Victor et al., 2018). The exacerbated production of pro-inflammatory mediators is well known for the deleterious effects, leading to chronic tissue injury (Nishida et al., 2017; Sousa-Victor et al., 2018).

What is the commonality of all these actions of CDNF and MANF? All the processes in which CDNF and MANF are involved, including neuro and cardioprotection, modulation of inflammation, homeostasis of pancreatic β-cells (Hummasti et al., 2010; Lindahl et al., 2014), among others, have ER-stress as their underlying cause. Interestingly, both proteins are ER-resident proteins, but differently from other ER/SR-resident stress-related proteins, their expression is enhanced upon ER/SR-induced stress, and more importantly they are secreted to the extracellular milieu to exert beneficial functions in neighboring cells. Thus, this new family of neuro/cardiotrophic factors has an intracellular activity, not yet completely understood, as well as extracellular functions that endow this new group of proteins with potential applications in medicine.

It has been shown that MANF expression and secretion also increase when rat cardiomyocytes (and HeLa cells) are treated with TG, but not with tunicamycin or DTT, other ER/SR-stressors that do not alter calcium homeostasis, leading to the proposition that MANF is retained in the ER/SR through a calcium-dependent mechanism, which relies on GRP78 forming a complex with MANF only in the presence of this ion (Glembotski et al., 2012). Once calcium concentration in the ER/SR is decreased (under stress conditions), the complex could dissociate leading to MANF secretion allowing its action as a cardiomyokine. Although MANF has a KDEL-like sequence (RTDL) in its C-terminal end, this degenerated sequence has a decreased affinity for the KDEL-R (Glembotski et al., 2012; Henderson et al., 2013) favoring its unbinding and secretion. The data presented here suggest that a similar mechanism of secretion might operate for CDNF in cardiomyocytes, although we did not investigate this deeply. Our main focus here was to discover how CDNF exerts its paracrine/autocrine functions in the cardiac context once the protein is secreted in response to an ER/SR insult.

Our data showed that heart hemodynamic function was significantly preserved upon CDNF treatment under pre- or post-conditioning regimens. We also showed that ventricular arrhythmias were significantly reduced by CDNF, as was infarct area (**Figure 4**). Interestingly, all these cardioprotective effects conferred by CDNF were abrogated when its interaction with the KDEL-R was blocked (**Figures 9 and 11**). We thus were able to identify the KDEL-R as a putative receptor for this neuro/cardiokine family of proteins, and PI3K/AKT as the signaling pathway that is activated upon receptor binding, at least for CDNF in the heart (**Figures 2, 4, 6 and 7**).

The KDEL receptor is a seven-transmembrane-domain protein whose function is to sort and retrieve proteins bearing a KDEL sequence (calnexin, GRP78, PDI, for instance) and possibly proteins with KDEL-like sequences (Erp72, MANF and CDNF) from the Golgi complex to the ER/SR (Dorner et al., 1990; Voutilainen et al., 2015; Becker et al., 2016). The importance of KDEL-R to cardiomyocyte homeostasis was demonstrated by studying transgenic mice that express a mutant KDEL-R in which reverse transport from Golgi to ER/SR was compromised. These animals die after birth with cardiac hypertrophy (Hamada et al., 2004). Recent studies, however, have suggested that the KDEL-R has additional functions and new cellular localizations, including the plasma membrane. In this regard, there is mounting evidence for the presence of KDEL-R at the surface of mammalian cells, where the receptor binds cargo proteins such as A/B microbial toxins K28, MANF and other proteins in which a KDEL-sequence is present at the C-terminus (Riffer et al., 2002; Henderson et al., 2013; Becker et al., 2016; Trychta et al., 2018). Trychta and collaborators (2018) have coined the term “ER exodosis” in an elegant study where they showed a massive departure of proteins containing KDEL-related sequences upon calcium-induced ER-stress in several mammalian cell lines. Even beneficial chaperones were among the proteins that were released.

The importance of the KDEL-R for MANF trafficking has been investigated in SH-SY5Y cells by using cells expressing different MANF constructs (with or without the RTDL sequence) in combination with different isoforms of the KDEL-R (KDEL-R1, 2 and 3) (Henderson et al., 2013). Interestingly, that study showed consistently that removal of the C-terminal RTDL sequence of MANF increased its secretion, reinforcing the idea that KDEL-R is an important partner of MANF in ER, although other studies questioned this interpretation (Norisada et al., 2016). Besides, the study conducted by Henderson and collaborators (Henderson et al., 2013) detected the presence of FLAG-tagged KDEL-R at the cell surface under ER-stress, and MANF binding to the cell surface required the RTDL sequence. This study unequivocally showed that KDEL-R located at the cell membrane modulates the binding of extracellular MANF, leading to the conclusion, as posed here, that the KDEL-R is the putative receptor for this family of NFs. Unfortunately, that study did not relate MANF-induced cardioprotection with its binding to cell membrane KDEL-R, a relation that we have shown here for CDNF, since the blockade of KDEL-R by peptides or the occlusion of KTEL sequence of CDNF by the antibody anti-CDNF abrogated the beneficial effects of CDNF on hearts and cardiomyocytes (**Figures 9 and 11**). Thus, the second novelty of the present study was to show that CDNF exerts its cardioprotective activity through binding to the KDEL-R that resides at the cell membrane, whose concentration increases upon ER/SR-stress (Becker et al., 2016). It is also important to note that the interaction with KDEL-R is pH dependent, being stronger at acidic pH (Wilson et al., 1993). Acidification occurs in ischemia, supporting the notion that MANF or CDNF interaction with KDEL-R at the cell membrane might be even stronger under ischemia.

For MANF, a recent study identified that the sulfoglycolipid sulfatide present in the outer cell-membrane leaflet plays a fundamental role in MANF internalization and cytoprotection in *C. elegans* and mammalian cells (Bai et al., 2018). CDNF did not require this sulfatide for its cellular internalization, but CDNF was not deeply explored in that study. Indeed, the N-domain of MANF and CDNF has a saposin-like fold, suggesting its direct interaction with membrane lipids without the necessity of a receptor. Our data, however, identified KDEL-R as a putative receptor for CDNF in the heart. How can we reconcile these observations? Calcium-induced ER/SR-stress increases the secretion of MANF and CDNF (and also other ER/SR-resident proteins), [**Figure 1** and (Bai et al., 2018)] to the extracellular milieu. ER-stress also enhances the expression of KDEL-R through the action of the transcription factor XBP1 (Trychta et al., 2018), increasing the presence of KDEL-R at the cell membrane (Henderson et al., 2013). Outside the cells, MANF and CDNF would establish an initial “unspecific” contact (mediated by sulfatide in the case of MANF) with the lipid bilayer through its N-domain and a further specific interaction with newly arrived KDEL-R through its C-terminal domain. Currently, we do not know whether the KDEL-R bound to CDNF is internalized and if so, what would be their fate. However, as presented here, CDNF/KDEL-R activates PI3K/AKT signaling. We do not know what is(are) the connector(s) between CDNF/KDEL-R and PI3K/AKT signaling, but we do know that the increase in p-AKT that takes place in hearts upon CDNF treatment is abrogated by THPKTEL, a peptide that binds to the KDEL-R at the cell membrane, but not by DRATSAL, a scrambled, ineffective peptide (**Figure 6D and Figure 10**).

There are several reports in the literature showing signaling pathways controlled by KDEL-R mainly in cancer or neuronal cells. Interestingly, GRP78, which is present at the cell membrane in several cancer cells, co-localizes with both subunits of PI3K, namely p85 and p110, as well as with PIP3 controlling PI3K/AKT pathway either directly or indirectly (Zhang et al., 2013). We are currently conducting additional experiments to dissect these missing connections.

Cardiac ischemia induces calcium transient depletion resulting in the halting of the contractile machinery and cell injury (Heusch, 2015). Immediate restoration of blood flow (reperfusion) is essential for myocardial survival, however, paradoxically, reperfusion itself can induces injury mediated by ER/SR stress, cytosolic calcium overload and mitochondrial impairment (Heusch, 2015). We have shown that CDNF prevents ER/SR-stress, by recovering calcium transient from cardiomyocytes. This effect would reduce calcium overload and prevent mitochondrial lesions and cell death (Heusch, 2015). Additionally, the treatment with CDNF prevents lesions by I/R. Similar results were observed with MANF (Glembotski et al., 2012), showing infarct size reduction in mice hearts. Indeed, CDNF reduces cerebral ischemic lesions by ER/SR-stress prevention (Zhang et al., 2018). Curiously, in our study, CDNF was able to prevent myocardial injury, not only in pre-ischemic treatment, but also in post-ischemic treatment. Since CDNF reduced reversible and irreversible I/R injuries, it could be a potential treatment against revascularization lesions. However, despite experimental studies having shown that interventions during myocardial reperfusion can reduce myocardial infarct size by up to 50%, clinical studies that use these strategies have not been successful (Yellon and Hausenloy, 2007). Thus, there is a necessity to develop new therapies against I/R injury. The Reperfusion Injury Salvage Kinase (RISK) pathway has emerged as a new target for preventing lethal reperfusion injury. There is a consensus, that the RISK pathway mediates the programmed cell survival, as well as an extensive preclinical evidence that activation of this pathway by pharmacological agents reduces the size of myocardial infarction by up to 50% (Hausenloy and Yellon, 2004). Cardioprotection by RISK pathway has been attributed to the activation of PI3K/AKT signaling pathway inducing inhibition of mitochondrial mPTP opening, sarcoplasmic reticulum Ca^2+^ uptake improvement, and antiapoptotic pathway activation (Yellon and Hausenloy, 2007; Heusch, 2015). These kinases are activated during myocardial reperfusion and confer cardioprotection avoiding lethal reperfusion injury (Hausenloy and Yellon, 2004). As shown in the present study, wortmannin, was the only tested inhibitor that was able to abrogate all the beneficial effects of CDNF. Altogether, these data unequivocally point to PI3K/AKT signaling as responsible for the cardiomyokine activity of CDNF. This was confirmed by increased levels of p-AKT after exogenous CDNF and return to basal levels in the presence of CDNF plus wortmannin. Interestingly, THPKTEL also reversed the increase in p-AKT levels induced by CDNF, connecting the binding of CDNF to the KDEL-R to PI3K/AKT activation.

In conclusion, the present study characterizes CDNF as new cardiomyokine acting at the cell membrane through its putative receptor KDEL-R and activating PI3K/AKT to exert its cardioprotective activity. All these findings are summarized as a model in **Figure 12**.

**Figure 12:**
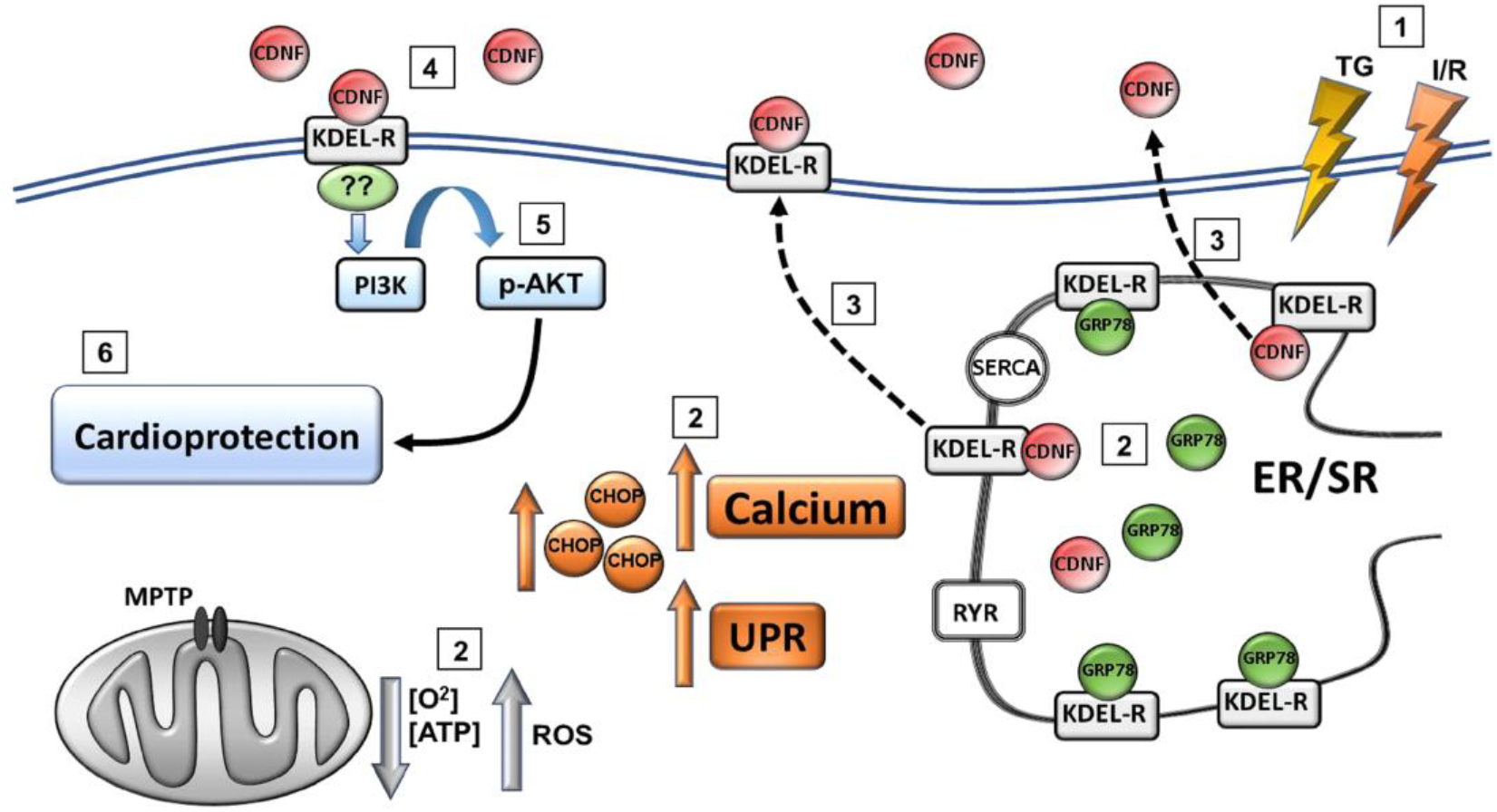
Schematic representation of the main findings of the present study. (**1**) TG or I/R induce cellular injuries causing (**2**) an increase in cytosolic calcium and UPR activation, up regulation of GRP78, CDNF and CHOP, as well as mitochondria impairment (decrease in [O2] and [ATP] and an increase in [ROS]). (**3**) CDNF is an ER/SR-resident protein remaining bound to KDEL-R through its degenerated KTEL sequence located at the C-terminal end. Under stress, KDEL-R and CDNF dissociate and migrate from the ER/SR to the cell membrane and extracellular milieu, respectively, (**4**) allowing the binding of CDNF to the KDEL-R at the cell surface with the subsequent (**5**) activation of PI3K/AKT signaling pathway either directly or indirectly conferring (**6**) cardioprotection to the cells. RYR= ryanodine receptor – releases calcium from the ER/SR; SERCA = sarco/endoplasmic reticulum Ca^2+^-ATPase – pumps calcium from the cytosol to the ER/SR; MPTP = mitochondrial permeability transition pore.

### Materials and Methods

All solutions were freshly prepared using MilliQ^®^ Water or high quality analytical grade organic solvents. All solutions were filtered before the use. The chemicals were obtained from Sigma-Aldrich (St. Louis, MO, USA) or purchased from vendors as indicated.

### CDNF production and purification

The plasmid pET-25b(+) containing a synthetic CDNF cDNA was transformed into Escherichia coli strain Rosetta Gami B (DE3) (Stratagene, San Diego, California, USA). Cell cultures were grown at 37°C and induced with 1mmol/L IPTG for 2h. The cells were harvested by centrifugation, resuspended in extraction buffer (20mmol/L MES at pH 6.0, Complete, EDTA-free Tabs—Roche, Basel, Switzerland) and lysed by sonication at 4°C. After centrifugation, the fraction containing the soluble proteins was loaded onto a 5mL Hitrap SP XL column (GE Healthcare, Chicago, Illinois, USA) equilibrated with 20mmol/L MES at pH 6.0. The bound material was eluted with a linear gradient of NaCl (0–1mmol/L) at 67mmol/L per min. Fractions containing the protein as checked by monitoring absorption at 280nm and denaturing polyacrylamide gel electrophoresis were loaded onto a Superdex 75 column (16 9 100 mm—GE Healthcare) equilibrated with 20mmol/L MES at pH 6.0 and 150mmol/L NaCl (see Latge et al., 2015 for more details).

### Animals

Adult male Wistar rats (300-350 g) and neonatal CD-1 mice (1-3 days old) were used following the Guide for the Care and Use of Laboratory Animals published by the US National Institutes of Health (8th edition, 2011) and the local Institutional Animal Care and Use Committee (100/16 and 015/17).

### Differentiation of human induced pluripotent stem cells into cardiomyocytes

Human induced pluripotent stem cells (hiPSCs) were differentiated into cardiac lineage following 30 days of induction as previously described (Lian et al., 2013). Briefly, 3×10^5^ hiPSCs were plated on 48 plate dish treated with 1% of Matrigel^®^ hESC-Qualified Matrix (Corning™, New York, USA) and cultivated on 2mTeSR™1 (STEMCELL Technologies, Vancouver, Canada) for 3 days. On day 0, the cells were cultivated in RPMI 1640 (Gibco™, Thermo Fisher, Waltham, Massachusetts, USA) supplemented with B-27™ (1640 (Gibco™) Supplement, without insulin (Gibco™) (RB-) and 9 μmol/L of CHIR 99021 (R&D SYSTEMS, Minneapolis, Minnesota, USA). After 24h, the cells were cultivated with RB-for the next 2 days and on day 3 and 4, the WNT signaling was inhibited with 10 and 5 μM of XAV939 (R&D SYSTEMS), respectively. On day 7, we observed the first beating areas and the cells were then cultivated in RPMI 1640 (Gibco™) supplemented with B-27™ Supplement (Gibco™) (RB+) until day 30^th^. These cells were called hiPSC-dCM (Human induced pluripotent stem cells differentiated into cardiomyocytes).

For efficiency evaluation of cardiomyocyte differentiation, after day 30, the hiPSC-dCM was dissociated using Tryple 1x (Gibco™) until the cells detached from the culture plate. Then the cells were centrifuged at 300g/5min. The cells were fixed with formaldehyde 4 % for 20 min at room temperature, permeabilized with PBS Triton 0.3 % for 30 min and stained with Troponin T (1:200) (Thermo Fisher) for 30 min AT 4°C. Afterwards the cells were stained with Alexa Fluor 647 goat anti mouse IgG (1:1,000) secondary antibody (Thermo Fisher) for 30 min at 4°C. The cells stained only with secondary antibody were used as fluorescence negative control. The data was acquired by BD Accuri™ C6 (BD Biosciences, San Jose, CA, USA) and analyzed with the FlowJo v10.1 software (FlowJo, USA).

### Isolation and culture of mice cardiomyocytes

The cardiomyocytes were isolated from hearts of neonatal mice (CD1-black) 1-3 days after birth. Briefly, the hearts were dissected out, placed in PBS and washed. The heart tissue was minced and digested in a dissociation solution (In mol/L: NaCl, 136.7; KCl, 2.68; Na_2_HPO_4_, 0.352; NaHCO_3_, 11.9; dextrose, 11) containing pancreatin (1.25mg/ml; Sigma-Aldrich) and bovine serum albumin (BSA; 3 mg/mL; Sigma-Aldrich) for 5 min at 37°C with gentle stirring. The supernatant fraction containing cells from each digestion was collected in a conical tube, suspended in growth medium containing 15% fetal bovine serum for inhibition of proteolytic enzymes and spun at 300 *g*/5 min. Elimination of non-muscle cells was achieved by pre-plating for 1h. After that, the cells were counted and seeded in 6-well plates at a density of 10^5^ cells/well and cultured in DMEM-high glucose containing 10% FBS at 37°C and 5 % CO_2_. Experimental in-vitro conditions were established 3 days after plating (Bagno et al., 2016).

### Immunocytochemistry imaging and quantification of fluorescence intensity

Cells were fixed with 4% paraformaldehyde for 10 min at room temperature. Then, cells were permeabilized in 0.3% triton X-100 in PBS and incubated with blocking solution (5% BSA, in PBS pH 7.4) for 1h. After incubation with appropriate primary (1:500), secondary antibodies (1:500) and Hoechst (1:5,000) cover slips were mounted with prolong. Images were acquired with an Olympus DS-Fi2 confocal microscope equipped with 63x oil immersion objective and with a Nikon DS-fi2 camera operated with the standard QC capture software (Leica). Quantification was performed with ImageJ software (NIH, Baltimore, MD) using the corrected total cell fluorescence (CTCF) method.

### Obtention of media and cell sample extracts

Cell media without fetal bovine serum were collected, appropriately concentrated, boiled and combined with the Laemmli sample buffer before SDS-PAGE and Western blot. The cells lysates were obtained by extracting cells in a minimal volume of cell lysis buffer composed of RIPA buffer (Sigma-Aldrich), EDTA 50 mmol/L (pH 8.0), sodium fluoride 5 mmol/L, 10% SDS, protease inhibitor mixture (Roche), phosphatase inhibitor mixture 1 (Sigma Aldrich) phosphatase inhibitor mixture (Sigma Aldrich). Samples of the media samples (150 μg) and cell extracts (50 μg) were analysed by SDS-PAGE, followed by Western blot.

### [Ca^2+^] Measurements

Human iPSC-dCM were plated on glass coverslips for 5 days and loaded with 5 μmol/L Fura-2AM at 37 °C in complete culture medium containing 2.5 mmol/L probenecid for 40-60 min before the intracellular calcium measurements. The cells were then accommodated in a chamber whose base was formed by the coverslip containing the cells that were maintained at 37 °C in the complete medium (volume of the incubation chamber – 500 μL). Cytoplasmic calcium concentrations of groups of hiPSC-dCM (20–30 cells) were measured on a fluorescence imaging spectrofluorimeter (Easy Ratio Pro equipped with a DeltaRAMX Illuminator, an Olympus IX71 microscope, a QuantEM 5125C camera and the Image-Pro Plus v 6.3 software; PTI Photon Technology International, Princeton, NJ). Fura-2 was excited alternately at 340 nm and 380 nm, and the emission was collected at 510 nm. The ratio measurement, which is proportional to the cytoplasmic calcium concentration, was determined every 100 ms for at least 5 min. Free intracellular Ca^2+^ concentration [Ca2+]i was monitored in arbitrary units as the F340/380 nm ratio. The amplitude of variations in [Ca^2+^]_i_ (ΔF340/F380) for cells without treatment and with treatment were obtained for 5 different intervals of 5-10 seconds from ratio measurements.

### Treatments applied to hiPSC-dCM and mouse cardiomyocytes before Western blot and calcium measurements

#### Conditions/Groups

<u>**Control**</u>: mouse cardiomyocytes or hiPSC-dCM cultures received no treatment before calcium measurement or Western blot analysis; <u>**Thapsigargin (TG)**:</u> mouse cardiomyocytes or hiPSC-dCM cultures were incubated with TG (1 μmol/L) during 20 h before calcium measurements or Western blot analyses; <u>**CDNF**:</u> mouse cardiomyocytes or hiPSC-dCM cultures were incubated with CDNF (1 μmol/L) during 20h before calcium measurements or Western blot analyses. <u>**TG+CDNF**:</u> hiPSC-dCM cultures were incubated with CDNF (1 μmol/L) one hour before the TG (1 μmol/L). The calcium measurements or Western blot analyses were performed after 20 h of incubation with TG. <u>**TG+CDNF+Wortmannin**:</u> hiPSC-dCM cultures were incubated first with wortmannin (0.3 μmol/L) for 15 min, then with CDNF (1 μmol/L for 1 h) and finally with TG (1 μmol/L). The calcium measurements were performed after 20h of incubation with TG. <u>**CDNF+Wortmannin**:</u> mouse cardiomyocytes or hiPSC-dCM cultures were incubated first with wortmannin (0.3 μmol/L) for 15min, then with CDNF (1μmol/L) for 1 h. The Western blot analyses were performed after 20 h of incubation. <u>**Wortmannin**:</u> hiPSC-dCM cultures were incubated with wortmannin (0.3 μmol/L) for 20h before the calcium measurements. The media was not changed after each substance addition in each group. The calcium measurements and/or Western blot analyses were performed after 20h of incubation in all groups.

### Langendorff experimental protocols and I/R

The I/R experiments were performed on isolated rat hearts as described previously (Maciel et al., 2017). The hearts were rapidly removed and cannulated through the aorta in a modified Langendorff apparatus and perfused at constant flow of 10 mL/min with Krebs-Henseleit buffer (KHB) solution: in mmol/L - NaCl 118; NaHCO_3_ 25; KCl 4.7; KH_2_PO_4_ 1.2; MgSO_4_ 1.2; CaCl_2_ 1.25, and glucose 11, at 37°C and equilibrated with a gas mixture of 95% O_2_ and 5% CO_2_ (pH 7.4). The perfusion temperature was held constant by a heat exchanger located next to the aortic cannula. A fluid-filled latex balloon was inserted through the left atrium into the left ventricle and connected to a pressure transducer and the PowerLab System (AD Instruments, Australia) for continuous left ventricular pressure recording. The left ventricular end-diastolic pressure (LVEDP) was set to 10mmHg by balloon inflation; during the experiment, the hearts were continuously immersed in 37°C warm buffer to avoid hypothermia. Hearts were allowed to stabilize for 20 min before a protocol was started. All hearts were submitted to 30min of basal recordings. The ischemia protocol was induced by full stop of retrograde perfusion during 30min. The reperfusion was performed by full re-establishment of retrograde perfusion during 10min for mitochondria analysis or 60min for infarct size measurements. *Langendorff experimental protocols:* **Conditions/Groups**: <u>**No-I/R**:</u> Hearts were perfused with KHB solution for 90min (only for mitochondria isolation); <u>**I/R**:</u> Global ischemia was induced for 30min by complete perfusion arrest followed by 10 or 60 min of reperfusion with KHB solution; <u>**CDNF-preconditioning**:</u> The hearts were perfused with the KHB containing CDNF (1 μmol/L) for 5min before 30min of global ischemia followed by 10 or 60 min of reperfusion with KHB. In the groups with antagonist, the substances were perfused for 5min before CDNF and along with CDNF (1 μmol/L) for 5min.; <u>**CDNF-postconditioning**:</u> Global ischemia was induced for 30 min by full stop of retrograde perfusion followed by the perfusion with KHB plus CDNF (1 μmol/L) during the first 5 min, followed by 5 or 55 min of reperfusion with KHB without CDNF. The antagonists used in the present study were: Wortmannin 0.3 μmol/L (PI3K-AKT inhibitor), Chelerythrine 10 μmol/L and Rottlerin 10 μmol/L (PKC inhibitors), and AG490 10 μmol/L (JAK-STAT3 inhibitor). The samples of isolated hearts were collected after 60min of reperfusion for Western-blot analyzes.

### Left ventricular developed pressure (LVDP), left ventricular end-diastolic pressure (LVEDP) and infarcted area measurements

LVDP and LVEDP were analysed at different time points during the experiments, namely, at baseline, every 5 min during ischemia, and every 5min during reperfusion until 60min of reperfusion. For this, the mean value of a 30 sec recording at the respective time point was taken. After 60 min reperfusion period, the hearts were sliced into 1.5 mm cross-sections from apex to base and incubated in 1% triphenyl tetrazolium chloride (TTC) for 4 min at 37 °C, followed by incubation in a 10% (v/v) formaldehyde solution for 24h to improve the contrast between the stained (viable) and unstained (necrotic) tissues. The infarct area was determined by planimetry using ImageJ software (version 1.22, National Institute of Health, USA). Infarct size was expressed as a percentage of the area at risk (total).

### ECG recordings in isolated hearts

Three silver-silver chloride electrodes were used to obtain electrocardiographic recordings. Two electrodes were connected to the differential input of a high-gain amplifier (BioAmp; AD Instruments) positioned close to the left ventricle and right atrium. The third electrode was grounded. The electrocardiograms were recorded with PowerLab/400 and Chart 4.0 software (AD Instruments).

### Peptides used in I/R experiments

The peptide sequences corresponding to the last 7 amino-acid residues of the C-terminal region from human (THPKTEL) and rat (TRPQTEL) CDNF or a scrambled peptide DRATSAL (Henderson et al., 2013) were synthesized by GenOne Biotechnologies with 98% purity. In the experiments where the peptides were tested separately, each peptide at 2 μmol/L was perfused during 5 min before I/R. When the peptides were tested in combination with CDNF, they were first perfused alone (2 μmol/L) and then together with CDNF (1 μmol/L) for 5 min before I/R

### Antibody used in I/R experiments

To reinforce the evidence that the KDEL receptor is involved in the protection induced by CDNF, 5 μg/ml of the antibody (Sigma Aldrich, St. Louis, MO) against peptides from the C-terminal domain of human CDNF were incubated with CDNF 1 μmol/L during 1 h before the experiment. After 1h of incubation, CDNF (1 μmol/L) plus the antibody (5μg/ml) were perfused for 5 min before the ischemia and reperfusion protocol.

### Western blots

SDS-PAGE gels were transferred to PVDF membranes. The membranes were probed with the following antibodies: CDNF (Sigma Aldrich; 1:1,000), GAPDH (Santa Cruz Biotechnology; 1: 10,000, Santa Cruz, CA, USA), GRP78 (Santa Cruz Biotechnology; 1:1,000), CHOP (Santa Cruz Biotechnology; 1:500), p-AKT (Cell Signalling; 1:750, Danvers, MA, USA), total AKT (Cell Signalling; 1:750), GRP78 (Sigma Aldrich; 1:1,000).

### Mitochondria isolation and measurements of mitochondrial function

Mitochondria isolation was performed according to (Gedik et al., 2017). After 10 min of reperfusion, the isolated hearts were rapidly removed, placed in ice-cold isolation buffer containing, in mmol/L: 250 sucrose, 10 HEPES, 1 ethylene glycol tetra acetic acid (EGTA), pH 7.4 with 0.5% w/v bovine serum albumin (BSA), minced thoroughly using scissors, and then homogenized with a tissue homogenizer (Ultra-Turrax) using two 10 sec treatments at a shaft rotation rate of 6,500 rpm to release subsarcolemmal mitochondria. This homogenate was further homogenized with proteinase type XXIV (8 IU/mg tissue weight, Sigma Aldrich) using a Teflon pestle for the release of the interfibrillar mitochondria from the tissue. The homogenate was centrifuged at 700g for 10 min at 4°C. The supernatant was collected and centrifuged at 12,000g for 10min. The resulting pellet was resuspended in isolation buffer without BSA and centrifuged at 10,000 g for min at 4°C. This procedure was repeated, and the pellet was resuspended in isolation buffer. The protein concentration of the isolated pellet was determined using a protein assay (Lowry method, Biorad, Hercules, CA, USA) by comparison to a BSA standard (Thermo Scientific, Waltham, MA, USA).

### Mitochondrial oxygen consumption measurements

Mitochondrial respiration was measured with a Clark-type electrode (Strathkelvin, Glasgow, UK) at 37°C during magnetic stirring in incubation buffer containing, in mmol/L: 125 KCl; 10 MOPS; 2 MgCl_2_; 5 KH_2_PO_4_; 0.2 EGTA with 5 pyruvate and 5 malate, as substrates for complex I. The oxygen electrode was calibrated using a solubility coefficient of 217 nmol O_2_/mL at 37°C. For the measurement of complex I respiration, mitochondria (corresponding to a mitochondrial protein amount of 100μg) were added to 1 mL incubation buffer. After 2 min of incubation, 1 mmol/L ADP was added and ADP-stimulated respiration measured over 2-3 min. Complex IV respiration was stimulated by adding N,N,N,N’-tetramethyl-p-phenylenediamine (TMPD, 300 μmol/L, Sigma Aldrich) plus ascorbate 3 μmol/L (Sigma Aldrich). Maximal uncoupled oxygen uptake was measured in the presence of 30 nmol/L carbonyl cyanide-p-trifluoromethoxyphenyl-hydrazone (FCCP, Sigma Aldrich) (Gedik et al., 2017).

### Mitochondrial oxygen consumption under simulated Hypoxia/Reoxygenation

To evaluate the direct effect of CDNF on isolated mitochondria, we measure the ADP-stimulated respiration after simulated hypoxia/reoxygenation with isolated mitochondria. The buffer (without pyruvate and malate) was made hypoxic by the introduction of purified nitrogen until the oxygen concentration was <15 nmol O_2_ mL^−1^. Mitochondria (200μg protein) were added to 0.5 mL hypoxic buffer without or supplemented with 1μmol/L CDNF. After 8 min, air-saturated incubation buffer (0.5 mL), again supplemented or not with 1 μmol/L CDNF, was added to achieve re-oxygenation for 2 min. After simulated hypoxia/reoxygenation or in the respective normoxic time control (10 min), pyruvate (5mmol/L) and malate (5mmol/L) were given as substrates for complex I; mitochondria were stimulated with ADP and respiration was measured over 2min. Complex IV respiration was stimulated by adding N,N,N,N’-tetramethyl-p-phenylenediamine (TMPD, 300μmol/L) plus ascorbate (3μmol/L). Maximal uncoupled oxygen uptake was measured in the presence of 30nmol/L carbonyl cyanide-p-trifluoromethoxyphenyl-hydrazone (FCCP) (Kleinbongard et al., 2015; Gedik et al., 2017).

### Mitochondrial ATP production measurements

After measurement of ADP-stimulated respiration, the incubation buffer containing mitochondria was taken from the respiration chamber and immediately supplemented with ATP assay mix (diluted 1:5, Sigma Aldrich)). Mitochondrial ATP production after each respiration measurement was determined immediately and compared with ATP standards using a 96-well white plate and a spectrofluorometer SpectraMax^®^ M3 (Molecular Devices California, EUA) at 560 nm emission (Gedik et al., 2017).

### Mitochondrial swelling and transmembrane potential measurements

The mitochondrial swelling and mitochondrial transmembrane potential were evaluated using a high resolution spectrofluorometer SpectraMax^®^ M3 (Molecular Devices California, EUA). The integrity of the mitochondrial membrane was assessed by varying the osmotic volumes of the mitochondria by spectrophotometric determination of the apparent absorption of the suspension at 540 nm. A mitochondrial suspension (100 μg/mL) was added to the breathing medium in the absence of respiratory substrates at 37 ° C and under constant stirring. The mitochondrial turgor was stimulated with 100 nmol/L of calcium. The swelling was expressed as a percentage of the absorption of the solution containing mitochondria in the presence of Ciclosporin A (0% swelling), in relation to the light emitted after the addition of FCCP 1μmol/L (100% swelling). For mitochondrial transmembrane potential (Δψ) determination, the probe TMRM (tetramethylrhodamine methyl ester, 400 nmol/L) was added to the incubator solution containing 100 μg/ml of mitochondria for 1h. The Δψ was estimated by the fluorescence emitted by TMRM under excitation of 580 nm. The Δψ was expressed as the percentage of fluorescence emitted by TMRM when mitochondria were incubated in the presence of cyclosporin A (0% Δψ), relative to the fluorescence emitted after addition of FCCP to fully depolarized mitochondria (100% Δψ).

### Extramitochondrial ROS concentration measurements

The Amplex Red Hydrogen Peroxide Assay (Life Technologies, Carlsbad, CA, USA) was used to determine extramitochondrial ROS concentration. Amplex Red reacts in 1:1 stoichiometry with peroxide in the presence of horseradish peroxidase (HRP) and produces highly fluorescent resorufin. The incubation buffer containing mitochondria was removed from the respiration chamber, and immediately supplemented with 50 μmol/L Amplex UltraRed and U/mL HRP. The supernatant was collected after 120 min of incubation in the dark. Extramitochondrial ROS concentration was determined and compared with H2O2 standards using a 96-well black plate and a spectrofluorometer SpectraMax^®^ M3 (Molecular Devices California, EUA) at 540 nm emission and 580 nm extinction wavelengths (Gedik et al., 2017).

### Statistics

Data are presented as mean ± standard error of the mean (S.E.M). The data were analysed by one-way ANOVA. When a significant difference was detected, one-way ANOVA was followed by Bonferroni post-hoc tests (GraphPad Prisma 6.0 software, San Diego, California, USA).

## Acknowledgements

This work was supported by Fundação Carlos Chagas Filho de Amparo à Pesquisa do Estado do Rio de Janeiro (FAPERJ) [E-26/010.003018/2014 and E-26/202.947/2017]; Coordenação de Aperfeiçoamento de Pessoal de Nível Superior (CAPES) [fellowships to the students]; and Conselho Nacional de Desenvolvimento Científico e Tecnológico (CNPq) [402967/2016-0]. We are grateful to Martha M. Sorenson for her suggestions and careful reading of the manuscript and Dr Diana Pellizari, Santiago Alonso and Elisabeth M. Duarte for lab technical assistance.

## Contributions

L.M., D.F.O, A.C.C.C., J.H.M.N. F.L.P, and D.F. Conception and design, Acquisition of data, Analysis and interpretation of data, Drafting or revising the article, Contributed unpublished essential data or reagents. L.O, F.M and H.A.S.S Acquisition of data, Analysis and interpretation of data. D.F. is the principal investigator. All authors discussed the results and commented on the manuscript.

